# A synthetic metabolic network for physicochemical homeostasis

**DOI:** 10.1101/698498

**Authors:** Tjeerd Pols, Hendrik R. Sikkema, Bauke F. Gaastra, Jacopo Frallicciardi, Wojciech M. Śmigiel, Shubham Singh, Bert Poolman

**Affiliations:** Department of Biochemistry, Groningen Biomolecular Sciences and Biotechnology Institute & Zernike Institute for Advanced Materials, University of Groningen, Nijenborgh 4, 9747 AG Groningen, The Netherlands

**Keywords:** Synthetic cell, membrane-bounded compartment, metabolic energy conservation, out-of-equilbrium network, physicochemical homeostasis, membrane transport

## Abstract

One of the grand challenges in chemistry is the construction of functional out-of-equilibrium networks, which are typical of living cells. Building such a system from molecular components requires control over the formation and degradation of the interacting chemicals and homeostasis of the internal physical-chemical conditions. The provision and consumption of ATP lies at the heart of this challenge. We report the *in vitro* construction in vesicles of a pathway for sustained ATP production that is maintained away from equilibrium by control of energy dissipation. We maintain a constant level of ATP with varying load on the system. The pathway enables us to control the transmembrane fluxes of osmolytes and to demonstrate basic physicochemical homeostasis. Our work demonstrates metabolic energy conservation and cell volume regulatory mechanisms in a cell-like system at a level of complexity minimally needed for life.

---

The generation and consumption of ATP lies at the heart of life. Complex networks of proteins, nucleic acids and small molecules sustain the essential processes of gene expression and cell division that characterize living cells, but without ATP they are non-functional. Herein lies one of the major challenges in the construction of synthetic cell-like systems. Other processes, such as achieving tunable DNA replication, efficient transcription and translation, and vesicle division (*1*) are essentially secondary to the solution of a controlled energy supply. Metabolic energy conservation is a prerequisite for synthetic systems no matter how complex. Energy is critical not just for (macro)molecular syntheses but also for maintaining the cytoplasm in a state compatible with metabolism through control over pH, ionic strength and solute composition. Here we have addressed that issue and show that we can control ATP production and ionic homeostasis in synthetic vesicles.

The bottom-up construction of synthetic cells from molecular components (*2*) differs in concept and strategy from the approach to engineer minimal cells, pioneered by the J. Craig Venter institute (*3*). Yet, both approaches address what tasks a living cell should minimally perform and how this can be accomplished with a minimal set of components. New biochemical functions and regulatory principles will be discovered as we make progress towards constructing a minimal cell. In the field of bottom-up synthetic biology (perhaps better called synthetic biochemistry), work is progressing towards establishing new information storage systems (*4*), replication of DNA by self-encoded proteins (*5*), the engineering of gene and protein networks (*6*), formation of skeletal-like networks (*7*), biosynthesis of lipids (*8,9*), division of vesicles (*10*) and non-lipid compartment systems (*11,12*). Protein synthesis using recombinant elements (*13*) incorporated into vesicles (*14,15*) or water-in-oil droplets (*12*) has been realized, but long-term sustained synthesis of chemicals is a bottleneck in the development and application of synthetic cell-like systems (*16,17*). At the root of the poor performance of reconstituted systems are challenges that relate to sustained production of nucleotides, import of substrate(s) and export of waste product(s), control of the internal physicochemical conditions (pH, ionic strength, crowding) and stability of the lipid-bounded compartment, all of which require constant energy dissipation.

Inspired by the challenges of the bottom-up construction of a living cell, we focus on the development of new open vesicle systems that sustain nucleotide levels and electrochemical gradients to allow further functionalities to be integrated. ATP is especially crucial, not only as a source of metabolic energy for most biological processes, but also as a hydrotrope, influencing the viscosity and possibly the structure of the cytosol (*18*). Energy consumption in a growing cell is dominated by polymerization reactions and maintenance processes (*19*), so regeneration of ATP is required to keep the cell running. Recent developments in the field of synthetic biochemistry have started to address the issue of ATP homeostasis. A cell-free molecular rheostat for control of ATP levels has been reported, employing two parallel pathways and regulation by free inorganic phosphate (*20*), but the system has not been implemented in vesicles. Photosynthetic artificial organelles have been constructed that form ATP on the outside of small vesicles, encapsulated in giant vesicles, allowing optical control of ATP dependent reactions (*21*).

Here, we present the construction of a molecular system integrated into a cell-like container with control of solute fluxes and tunable supply of energy to fuel ATP-requiring processes. We have equipped the vesicles with sensors for online readout of the internal ATP/ADP ratio and pH, allowing us to conclude that the system enables long-term metabolic energy conservation and physicochemical homeostasis.

## RESULTS AND DISCUSSION

### A system for sustained production of ATP

The conversion of arginine into ornithine, ammonia plus carbon dioxide yields ATP in three enzymatic steps (Fig. 1A) (*22*). For sustainable energy conservation in a compartmentalized system, the import of substrates and excretion of products have to be efficient, which can be achieved by coupling the solute fluxes. The antiporter ArcD2 facilitates the stoichiometric exchange of the substrate arginine for the product ornithine (*23*), which is important for maintaining the metabolic network away from equilibrium. The thermodynamics of the arginine conversion under standard conditions are given in Fig. 1B. The equilibrium constant of the conversion of citrulline plus phosphate into ornithine plus carbamoyl-phosphate is highly unfavorable, but the overall standard Gibbs free energy difference (ΔG^o^) of the breakdown of arginine is negative. Since the actual ΔG is determined by ΔG^0^ and the concentration of the reactants, the antiport reaction favors an even more negative ΔG by maintaining an out-to-in gradient of arginine and in-to-out gradient of ornithine (Fig. 1C). We anticipate that NH_3_ and CO_2_ will passively diffuse out of the cell.

**Figure 1.**
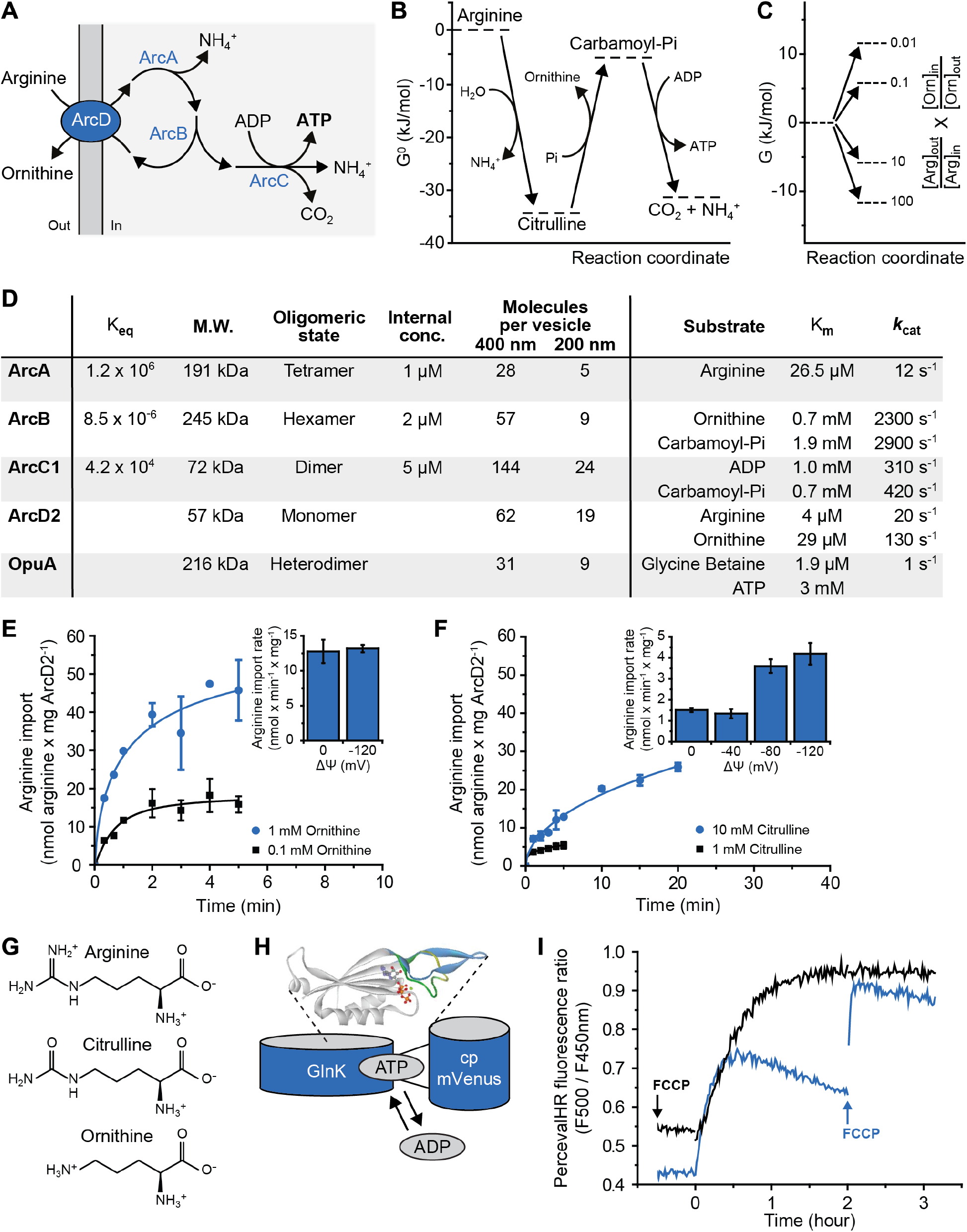
A system for sustained production of ATP. (**A**) Schematic representation of the arginine breakdown pathway. (**B**) Thermodynamics of arginine conversion for the reactions of ArcA, ArcB and ArcC1. G^0^ values were calculated for pH 7.0 and an ionic strength of 0.1 M using eQuilibrator 2.2. (**C**) Thermodynamics of the arginine transport reaction. G values were calculated at varying concentration gradients of arginine (outside to inside) and ornithine (inside to outside). (**D**) Molecular and kinetic properties of the enzymes; K_eq_ was calculated as in (B). The K_M_ values of a given substrate were determined under conditions of excess of the other substrate. The number of molecules per vesicle was calculated from the internal concentration of the enzymes and the average size of the vesicles, for 400 nm (left column) and 200 nm (right column) extruded vesicles (see Fig. S1) Kinetic parameters of ArcB are given for the back reaction. The kinetic parameters of ArcD2 were estimated from measurements in cells (*35*), assuming that ArcD2 constitutes 1% of membrane protein; the data for OpuA are from (*33*). (**E**) Kinetics of arginine uptake in proteoliposomes with 1 mM (blue circles) and 0.1 mM (black squares) ornithine on the inside (*Protocol B2*); ^14^C-arginine concentration of 10 µM. Inset: influence of a membrane potential (ΔΨ) on arginine uptake with 1 mM ornithine on the inside. Data from replicate experiments (*n*=2) are shown. (**F**) Kinetics of arginine uptake in proteoliposomes with 10 mM (blue circles) and 1 mM (black squares) citrulline on the inside (*Protocol B2*); ^14^C-arginine concentration of 10 µM. Inset: influence of a membrane potential (ΔΨ) on arginine uptake with 10 mM citrulline on the inside (*n*=2). (**G**) Structures of arginine, citrulline and ornithine at neutral pH. (**H**) Schematic representation of PercevalHR; adapted from *(25)*. (**I**) Effect of FCCP on the fluorescence readout of PercevalHR (*Protocol A1*); the ratio of the fluorescence peaks at 500 nm and 450 nm is shown. In the absence of FCCP, the fluorescence readout declines 30 min after addition of 10 mM arginine (at t=0) due to changes in the internal pH (addition of FCCP after 2 hours increases the signal, indicated by the blue arrow). The fluorescence of signal is constant for several hours in the presence of 10 µM FCCP (*n*=2).

To construct the system for ATP regeneration, we purified and characterized arginine deiminase (ArcA), ornithine transcarbamoylase (ArcB), carbamate kinase (ArcC1), and the antiporter ArcD2. Their kinetic and molecular properties are summarized in Fig. 1D. The enzymatic network is enclosed with inorganic phosphate and Mg-ADP in vesicles composed of synthetic lipids, while ArcD2 is reconstituted in the membrane. The concentration and number of reporters, ions and metabolites per vesicle is given in **Supplementary Table 4**. The lipid composition of the vesicles is based on general requirements for membrane transport (bilayer and non-bilayer-forming lipids, anionic and zwitterionic lipids), which is tuned to our needs (*see below*; *24*). The vesicles obtained by extrusion through 400 and 200 nm filters have an average radius of 84 nm (SD=59 nm; n=2090) and 64 nm (SD=39 nm; n=2092) respectively, as estimated from cryo-electron microscopy images; the average internal volume of the vesicles centers around radii of 226 nm (SD=113 nm) and 123 nm (SD=49 nm), respectively (**Supplementary Fig. 1**). Although a fraction of the vesicles is multi-lamellar, it is likely that all layers of the vesicles are active because we reconstitute the membrane proteins in liposomes prior to the inclusion of the enzymes, sensors and metabolites (**Supplementary methods**, *protocol A1*). The encapsulation of the luminal components is done by five freeze-thaw cycles, which induce content exchange between vesicles and homogenize the membranes. The vesicles obtained by extrusion through 200 nm filters are more homogenous in size (**Supplementary Fig. 1**) but contain a smaller number of components (Fig. 1D; **Supplementary Table 4**), yet the performance of the metabolic network is similar in both types of compartments (*see below*).

We first characterized ArcD2 in lipid vesicles without the enzymatic network and demonstrated exchange of arginine for ornithine (Fig. 1E). ArcD2 transports arginine *in* and ornithine *out* in both membrane orientations, which is a property of this type of secondary transporter. The direction of transport is determined by the concentration gradients of the amino acids, not by the orientation of the protein. The arginine/ornithine antiport reaction is not affected by an imposed membrane potential (ΔΨ) (Fig. 1E, **inset**). Surprisingly, we also detect that ArcD2 exchanges arginine for citrulline, albeit at a much lower rate than the arginine/ornithine antiport (Fig. 1F). The arginine/citrulline antiport is electrogenic (Fig. 1F, **inset**), which agrees with arginine (and ornithine) being cationic and citrulline being neutral at pH 7 (Fig. 1G).

The turnover number (*k*_*cat*_) and equilibrium constant (K_eq_) of the enzymes were used to guide the initial design of the pathway, and the enzymes were incorporated in the vesicles at a copy number well above the stochastic threshold (Fig. 1D). The ArcD2 protein is reconstituted at a lipid-to-protein ratio of 400:1 (w/w), yielding on average 62 and 19 antiporters per vesicle with radii of 226 and 123 nm, respectively. Since arginine is imported when a counter solute is present on the inside, we include L-ornithine in the vesicles to enable the metabolism of arginine. For readout of ATP production, we enclosed PercevalHR (*25*), a protein-based fluorescent reporter of the ATP/ADP ratio (Fig. 1H); the calibration and characterization of the sensor are shown in **Supplementary Fig. 2**. Upon addition of arginine, the vesicles produce ATP. Thus, after an initial, rapid, increase in the ratio of the excitation maxima at 500 nm and 430 nm (representing an increase in ATP/ADP ratio) the ratio unexpectedly declines after 30 min (Fig. 1I, **blue trace**). The ATP/ADP ratio increases again and stabilizes (Fig. 1I, **black trace**) in the presence of the protonophore carbonyl cyanide-4-(trifluoromethoxy) phenylhydrazone (FCCP), suggesting that arginine conversion by the metabolic network changes the internal pH of the vesicles (*see below*). The drop in fluorescence signal without FCCP is explained by the pH-dependent binding of nucleotides to PercevalHR (**Supplementary Fig. 2B and 2C**).

We found that, in the initial design of the pathway, a substantial amount of citrulline is formed on the outside, which is due to residual binding of ArcA to the outer membrane surface even after repeated washing of the vesicles. We thus inactivated external ArcA by treatment of the vesicles with the membrane-impermeable sulfhydryl reagent *p*-chloromercuribenzene sulfonate (pCMBS). To avoid inhibition of the antiporter ArcD2, we engineered a cysteine-less variant that is fully functional and insensitive to sulfhydryl reagents (**Supplementary Fig. 3**). This optimized system is used for further characterization and application.

### Arginine breakdown and control of futile hydrolysis and pH

The breakdown of arginine, in the lumen of the vesicle, is given by the reaction equation:

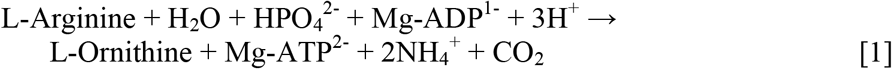

The external pH increases upon the addition of arginine (Fig. 2A), which is in accordance with Eq. 1 when the reaction products (except for ATP) end up in the outside medium (Fig. 2B). Unexpectedly, the vesicle lumen acidifies over longer timescales, that is, after an initial increase of the internal pH (Fig. 2C, **blue line**; pyranine calibration shown in **Supplementary Fig. 4**); the transient in the internal pH is more evident when the internal buffer capacity is decreased (Fig. 2C, **black line**). The acidification of the vesicle lumen cannot be readily explained if arginine is solely converted into ornithine (Eq. 1). Indeed, we found citrulline as an end product in addition to ornithine (Fig. 2D). This side reaction is possible if steps in the pathway downstream of ArcA are limiting the breakdown of arginine (Fig. 2E, **bold arrow**) and when citrulline is exchanged for arginine (Fig. 2E, **dashed arrow**). We present a more detailed explanation for the futile hydrolysis of arginine in **Supplementary Text S1**.

**Figure 2.**
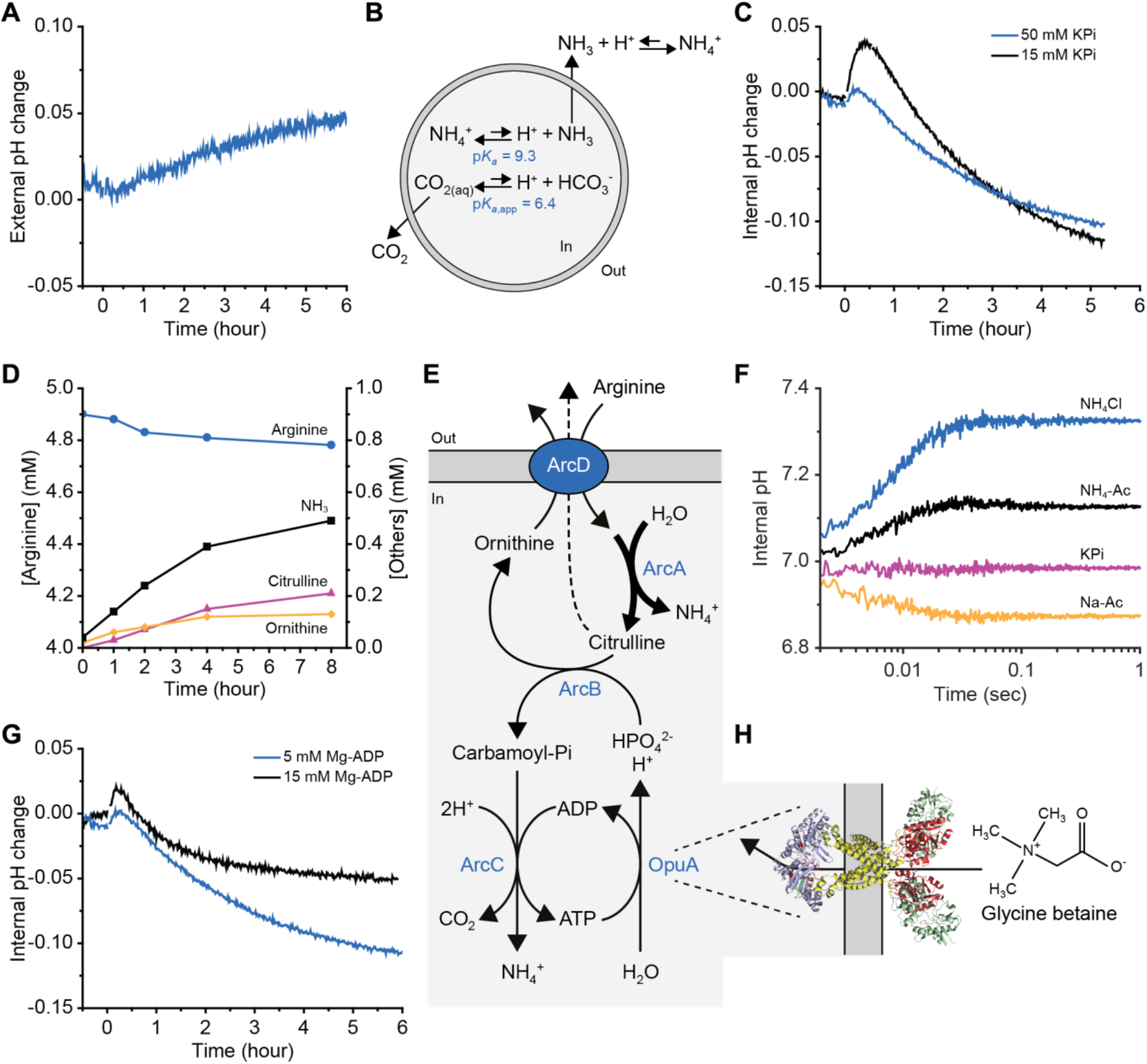
Arginine breakdown and control of futile hydrolysis and pH. The internal composition of the vesicles at the start of the experiment is given in Fig. 1D (enzymes) and **Supplementary Table 4** (ions, metabolites) and the preparation of vesicles is given in *Protocol A1*, unless specified otherwise. (**A**) External pH change (*Protocol B5)* of arginine metabolizing vesicles in outside medium with 10 mM KPi pH 7.0 plus 355 mM KCl. The ATP production was started by adding 5 mM arginine at t=0. (**B**) Schematic representation of the pH effects caused by ammonia and carbon dioxide diffusion. (**C**) Internal pH change (*Protocol B4*) of arginine metabolizing vesicles with either 50 mM (blue trace) or 15 mM KPi plus 40 mM KCl, pH 7.0 on the inside (black trace, *Protocol A2*); 5 mM arginine was added at t=0 (*n*=2). (**D**) External concentration of metabolites (*Protocol B6*): arginine (blue circles), citrulline (pink triangles), ornithine (yellow diamonds) and NH_3_ (black squares), as measured by HPLC in vesicles treated with 25 µM pCMBS; 5 mM arginine was added at t=0. (**E**) Schematic representation of arginine breakdown; the futile hydrolysis of arginine and arginine/citrulline exchange are depicted by bold and dashed arrows, respectively. (**F**) Stopped-flow fluorescence measurements to determine the permeability of the vesicles for NH_4_Cl (blue trace), NH_4_-acetate (black trace), potassium phosphate (pink trace) and sodium acetate (yellow trace); pyranine inside the vesicles was used as pH indicator (*n*≥2). (**G**) Internal pH change (*Protocol B4*) of arginine metabolizing vesicles with either 5 mM Mg-ADP (*Protocol A1*, blue trace) or 15 mM Mg-ADP on the inside (*Protocol A3*, black trace); 5 mM arginine was added at t=0 (*n*=2). (**H**) Homology model of OpuA and structure of the compatible solute glycine betaine. Glycine betaine import via OpuA consumes the ATP as indicated in Fig. 2E.

How do the side reactions of the arginine breakdown pathway lead to acidification of the vesicle lumen? The pK_A_ of NH_4_^+^ ↔ NH_3_ + H^+^ is 9.09 at 30 °C (*26*), and thus at pH 7.0 the fraction of NH_3_ is small, but the base/conjugated acid reaction is fast. If NH_3_ diffuses across the membrane, it will leave a proton behind in the vesicle lumen. Since the external volume is large compared to the internal one, there will be a net flux of NH_3_ from the vesicle lumen to the medium. Using stopped-flow fluorescence-based flux measurements to probe the permeability of the vesicles for small molecules, we confirmed that NH_3_ diffuses out rapidly, but that the membrane is highly impermeable for inorganic phosphate, K^+^, and Cl^−^ ions (Fig. 2F). CO_2_ also diffuses rapidly across the membrane, down its concentration gradient, but only NH_3_ leaves behind protons, therefore it is this that causes the pH change in the vesicle lumen. Finally, the breakdown of arginine to ornithine plus NH_4_^+^ and CO_2_ is a dead-end process, which reaches equilibrium if the produced ATP is not utilized; the system runs out of ADP in about 30 min. The production of NH_4_^+^ from the conversion of arginine to citrulline then takes over, and the accompanying diffusion of NH_3_ out of the vesicles leads to a net acidification of the vesicle lumen (Fig. 2C). Indeed, Fig. 2G shows that the vesicles acidify significantly less when the vesicles are loaded with a higher concentration of ADP and the ATP synthesis is extended.

### Physicochemical homeostasis

Cell growth is impacted by the solute concentration of the environment. Control of osmolyte import and export under conditions of osmotic stress allows cells to maintain their volume and achieve physicochemical homeostasis (*27–29*). Potassium is the most abundant osmolyte in many (micro)organisms, but excessive salt accumulation increases the ionic strength, which diminishes enzyme function. To control the volume, internal pH, and ionic strength, bacteria modulate the intake of potassium ions. When needed, they replace the electrolyte for so-called compatible solutes, like glycine betaine, proline and/or sugars (*30*). Compatible solutes like glycine betaine not only act in volume regulation but also prevent aggregation of macromolecules by affecting protein folding and stability (*31,32*).

The energy produced by the ATP breakdown pathway has been used to modulate the balance of osmolytes in the vesicles via activating the ATP-driven glycine betaine transporter OpuA (Fig. 2H). To this end we co-reconstituted OpuA with the components of the metabolic network for ATP production. OpuA transports solutes into the vesicle lumen when the protein is oriented with the nucleotide-binding domains on the inside. It happens to be that we reconstitute OpuA for more than 90% in this desired orientation (*24,33*), but any protein in the opposite orientation is non-functional because ATP is only produced inside the vesicles. In vesicles with 13 mole% DOPG [1,2-di-(9Z-octadecenoyl)-*sn*-glycero-3-phospho-(1’-*rac*-glycerol)] OpuA is constitutively active and imports glycine betaine at the expense of ATP (Fig. 3A), albeit at a low rate (*34*). OpuA is ionic strength-gated and more active when sufficient levels of anionic lipids are present in the membrane (*24,34*), which is the physiologically more relevant situation; for this, we use 38 mole% DOPG and an internal ionic strength below 0.2 M to lock OpuA in the off state. By increasing the medium osmolality with membrane-impermeant osmolytes (KPi or KCl), the vesicles shrink due to water efflux (Fig. 3B) until the internal osmotic balance is achieved, which occurs on the timescale of less than 1s. Thus, the pressure exerted by the addition of KPi or KCl is dissipated by lowering the volume-to-surface ratio of the vesicles. The accompanying increase in internal ionic strength activates OpuA and glycine betaine is imported to high levels (Fig. 3C, **blue circles**). The vesicles now possess an interior that is a mixture of glycine betaine and salts. The consumption of ATP by the gated import of glycine betaine is shown in Fig. 3D. Most remarkably, the gated import continues for hours when the internal ionic strength remains above the gating threshold and the pH is kept constant (Fig. 3C, **blue circles**). The open system, with arginine feed and product drain, performs at least an order of magnitude better than closed systems for ATP regeneration, as exemplified by the creatine-phosphate/kinase system (Fig. 3C, **pink triangles**) (*33*). Comparable results were obtained when smaller, yet more homogenous vesicles were formed by extrusion through 200 nm polycarbonate filters (Fig. 3C, **black squares**, see also **Supplementary Fig. 5**). We calculate that at least 70% of the vesicles with OpuA are fully functional in metabolic energy conservation and homeostasis (**Supplementary Text S2**).

**Figure 3.**
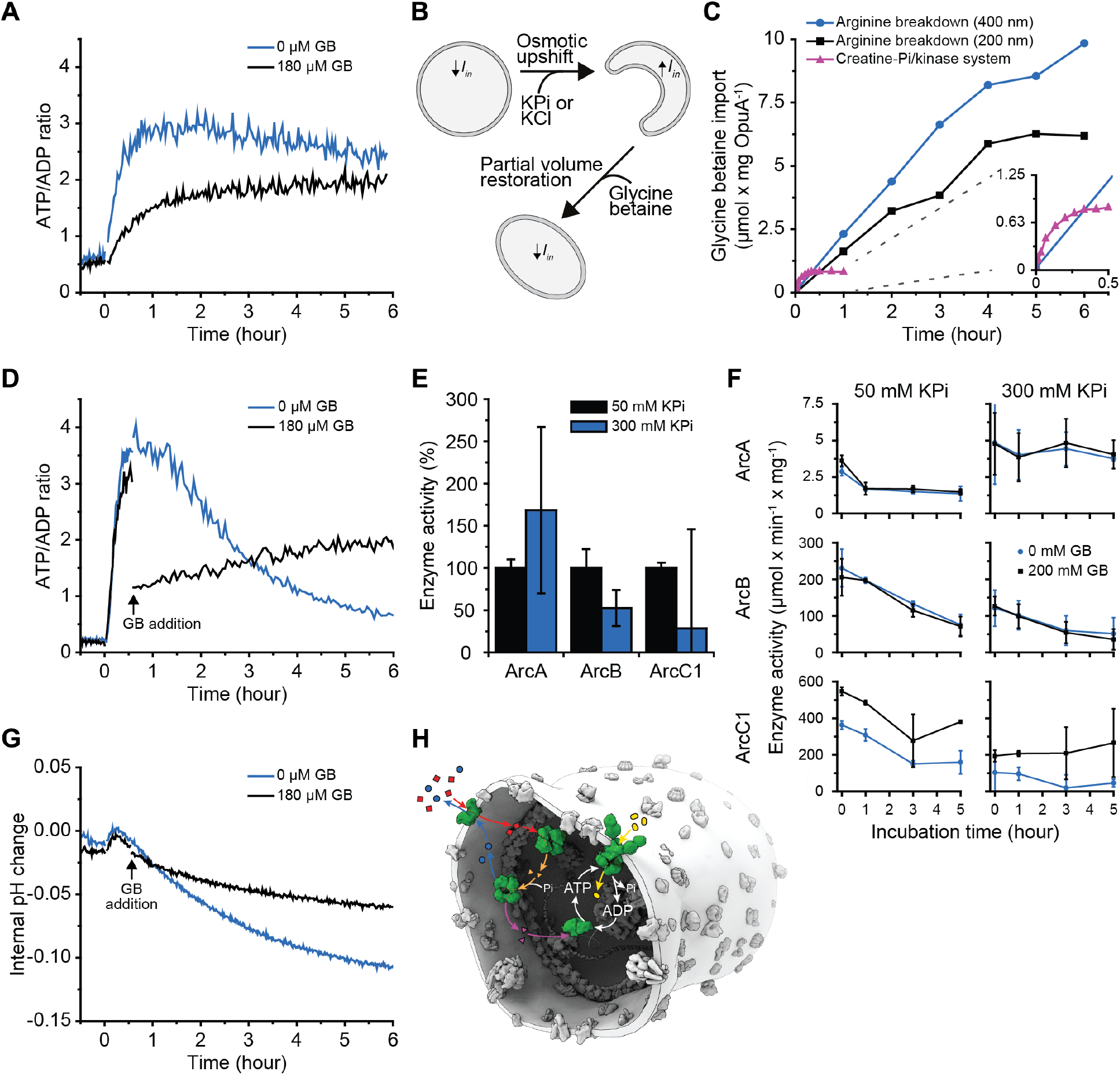
Physicochemical homeostasis of arginine-metabolizing vesicles. The internal composition of the vesicles at the start of the experiment is given in Fig. 1D (enzymes) and **Supplementary Table 4** (ions, metabolites) and the preparation of vesicles is given in *Protocol A1* (with 38 mole% of DOPG), unless specified otherwise. (**A**) Effect of external glycine betaine (GB) on the ATP/ADP ratio measured by PercevalHR (*Protocol B3*) in unshocked arginine-metabolizing vesicles (5 mM added at t=0) in the presence (black trace) and absence (blue trace) of 180 µM glycine betaine, added at t= −0.5 h (*n*=2); the vesicles were made with 13 mole% of DOPG. (**B**) Schematic representation of the effect of osmotic upshift and partial volume restoration through glycine betaine uptake are shown. (**C**) Comparison of glycine betaine uptake (*Protocol B1*) driven by ATP formed in the arginine breakdown pathway (blue circles: 400 nm extruded vesicles; black squares: 200 nm extruded vesicles, *Protocol A4*) and the creatine-phosphate/kinase system (pink triangles, *Protocol A5*) (*n*=2). (**D**) Effect of glycine betaine (GB) on the ATP/ADP ratio measured by PercevalHR (*Protocol B3*) inside arginine-metabolizing vesicles exposed to an osmotic upshift (addition of 250 mM KCl externally) in the presence (black trace) and absence (blue trace) of 180 µM glycine betaine (added at t=0.5 h); 5 mM arginine was added at t=0 (*n*=3). (**E**) Activity of ArcA (left), ArcB (middle) and ArcC1 (right) in 50 mM KPi, pH 7.0 (black bar) and 300 mM KPi, pH 7.0 (blue bar) as determined from the production of citrulline (ArcA, ArcB) or ATP (ArcC1). The activities were normalized to those in 50 mM KPi, pH 7.0; the absolute activities are given in **Supplementary Table 5** (*n*=2). (**F**) Stability of ArcA (top), ArcB (middle) and ArcC1 (bottom) in 50 mM KPi, pH 7.0 (left) and 300 mM KPi, pH 7.0 (right) at 30 °C, in the presence (black squares) and absence (blue circles) of 200 mM glycine betaine (*n*=2). (**G**) Effect of glycine betaine on the pH measured by pyranine (*Protocol B4*) inside arginine metabolizing vesicles exposed to an osmotic upshift (250 mM KCl) in the presence (black trace) and absence (blue trace) of 180 µM glycine betaine (added at t=0.5 h); 5 mM arginine was added at t=0 (*n*=3). (**H**) Schematic representation of the synthetic metabolic network in the cell-like container (arginine, red squares; ornithine, blue circles; citrulline, orange triangles; carbamoyl-phosphate, pink triangles; glycine betaine, yellow ovals; NH_3_ and CO_2_ are not shown).

Next, we tested how the physicochemical homeostasis is sustained when the vesicles are osmotically challenged. Fig. 3D shows the evolution in time of the ATP/ADP ratio upon addition of arginine to vesicles that were exposed to an increased medium osmolality 30 min before the addition of arginine; FCCP was added to avoid pH effects on the readout of PercevalHR (**Supplementary Fig. 2B**). In the absence of glycine betaine the ATP/ADP ratio peaks at 1 hour and decreases over the next 4 to 5 hours (Fig. 3D, **blue line**), which indicates the presence of futile ATP hydrolysis when the internal salt concentration is high. Intriguingly, when glycine betaine was added 0.5 hour after arginine (Fig. 3D, **black line**), the ATP/ADP ratio drops instantly, but then remains stable for 6 hours. Glycine betaine therefore has two effects: its accumulation provides a ‘cytosol’ that is more compatible with enzyme function (Fig. 3E; F), but it also provides a metabolic sink for ATP through its OpuA-mediated transport. The decrease in ATP/ADP ratio in the absence of glycine betaine can be explained by inhibition of enzymatic activity at high ionic strength (Fig. 3E), combined with futile hydrolysis of ATP. Accordingly, the decrease in ATP/ADP ratio is much less in vesicles in which the ionic strength is kept low (Fig. 3A). Finally, we report ATP/ADP ratios because the absolute concentrations change when the vesicles are osmotically shrunk and subsequently regain volume. In most experiments, the initial adenine nucleotide (=ADP) concentration was 5 mM but increases when the vesicles are exposed to osmotic stress. In Fig. 3A we report data of unshocked vesicles and here the ATP/ADP ratios of 2 to 3 correspond to 3.33 to 3.75 mM of ATP, which is in the range of concentrations in living cells.

Importantly, the introduction of an ATP-consuming process stimulates the full pathway at the expense of citrulline formation, and hence should stabilize the internal pH. Indeed, our results show that the system is better capable of maintaining the internal pH relatively constant when ATP is utilized (Fig. 3G, **black line**). Under these conditions the arginine-to-citrulline conversion is diminished relative to full pathway activity. We therefore propose that the stabilizing effect of glycine betaine accumulation originates from a lowering of the ionic strength (partial restoration of the vesicle volume), a chaperoning effect on the proteins and the maintenance of the internal pH, hence reflecting what compatible solutes do in living cells (*30*).

## Conclusions

We present the *in vitro* construction of a cell-like system that maintains a metabolic state far-from-equilibrium for many hours (Fig. 3H). This is one of the most advanced functional reconstitutions of a chemically defined network ever achieved, which allows the development of complex life-like systems with adaptive behavior in terms of lipid and protein synthesis, cell growth and intercellular communication. We show that ATP is used to fuel the gated transport of glycine betaine, which allows the synthetic vesicles to maintain a basic level of physicochemical homeostasis. Maintenance of the ATP/ADP ratio, internal pH, and presumably ionic strength are crucial for any metabolic system in the emerging field of synthetic biochemistry (*17,20*). We expect that our metabolic network will find wide use beyond membrane and synthetic biology, as biomolecular out-of-equilibrium systems will impact the development of next generation materials (*e.g.* delivery systems) with active, adaptive, autonomous and intelligent behavior.

## Supporting information

Supplemental Methods and Data for SynCell paper 10.1

## Acknowledgments

We thank Giorgos Gouridis for the construction of the pNZarcA and pNZarcB vectors, Marc Stuart for assistance with the cryo-EM measurements, Cecile Deelman-Driessen for enzyme assays, Michiel Punter for assistance with data fitting, Dirk-Jan Slotboom and Juke Lolkema for discussions, and Wilhelm Huck and Ian Booth for critical reading of the manuscript.

## Author contributions

BG, BP, HS, TP and WS designed the research; BG, HS, TP and WS performed most of the research; SS purified enzymes; JF performed the stopped-flow measurements; BG, BP, HS, TP and WS analyzed data; and BG, BP, HS and TP wrote the paper.

## Data and materials availability

All data is available in the main text or supplementary materials. All data, code, and materials used in the analysis are available to any researcher.

## Supplementary Materials and Data

**Supplementary Text, text boxes S1 and S2**

**Materials and methods**, including Supplementary Tables 1 to 4; for experimental details we refer to Protocols A1 to A6 (vesicle preparations) and B1 to B7 (assays).

**Supplementary data**

Supplementary Figures 1 to 7

Supplementary Table 5

**Supplementary references** (36 - 49)

