## Supplemental Methods and Data for SynCell paper 10.1 for "A synthetic metabolic network for physicochemical homeostasis"

**This PDF file includes:**

**Supplementary Text, text boxes S1 and S2**
**Materials and methods**, including Supplementary Tables 1 to 4; for experimental details we refer to Protocols A1 to A6 (vesicle preparations) and B1 to B7 (assays).
**Supplementary data**
Supplementary Figures 1 to 7
Supplementary Table 5
**Supplementary references** (36 - 49)

### SUPPLEMENTARY TEXT

**Text S1. Citrulline as byproduct and arginine/citrulline antiport**

Since external ArcA was inactivated by pCMBS, we infer that part of the citrulline is not metabolized further but exported and accompanied by diffusion of $NH_3$ through the membrane. The production of citrulline is possible if steps in the pathway downstream of ArcA are limiting the breakdown of arginine and when citrulline is exchanged for arginine (Fig. 2E, dashed arrow). In the vesicles with the full arginine breakdown pathway, citrulline will compete with ornithine for export, when citrulline internal concentration is high. The equilibrium constant of the reaction catalyzed by ArcB ($K_{eq} = 8.5 \times 10^{-6}$, see Fig. 1D) predicts high citrulline-to-ornithine ratios in the vesicles. Indeed, we find that the internal citrulline-to-ornithine ratio increases from 0 to more than 10 when arginine is converted for 1h. Thus, the $K_{eq}$ values of the reactions of citrulline formation and breakdown (Fig. 1D), and the substrate promiscuity of ArcD2 (Fig. 1E; F) enable arginine/citrulline in addition to arginine/ornithine antiport.

**Text S2. Analysis of fraction of vesicles with full ATP breakdown pathway and ATP consumption**

To determine the fraction of vesicles with a fully functional arginine breakdown pathway we compared the rates of transport of glycine betaine in our synthetic cell system with those of OpuA vesicles containing 10 mM of ATP, e.g. Fig. 3B. Transport only occurs when ATP is formed, which requires the presence of each of the enzymes well above the stochastic threshold.

When the metabolic pathway reaches steady state, the ATP and ADP concentrations are about 3.3 and 1.7 mM, respectively (See Table S4, ATP/ADP ratio of 2). The $K_M$ value for ATP is 3 mM and the $K_I$ for ADP is 12 mM (taken from *36*). From these numbers we compute the $V/V_{MAX}$ for transport in the synthetic vesicles. For the OpuA vesicles with 10 mM ATP there is negligible formation of ADP when initial rates of transport are determined, and the $V/V_{MAX}$ is calculated similarly. From the ratio of the V/Vmax in the vesicles with full pathway over the $V/V_{MAX}$ in the OpuA vesicles (data of Fig. 3B and comparable experiments), we obtain a lower limit for the fraction of active vesicles of 70%.

### MATERIALS AND METHODS

#### Materials

Common chemicals were of analytical grade and ordered from Sigma-Aldrich Corporation, Carl Roth GmbH & Co. KG or Merck KGaA. The lipids were obtained from Avanti Polar Lipids, Inc. (>99 % pure, in chloroform): 1,2-dioleoyl-*sn*-glycero-3-phosphoethanolamine (DOPE) [850725C], 1,2-dioleoyl-*sn*-glycero-3-phosphocholine (DOPC) [850375C] and 1,2-dioleoyl-*sn*-glycero-3-phospho-(1'-rac-glycerol) (DOPG) [840475C]. *n*-dodecyl-β-D-maltoside (DDM) [D97002] was purchased from Glycon Biochemicals GmbH and Triton X-100 [T9284] from Sigma-Aldrich Corporation. $^{14}$C-glycine betaine was prepared enzymatically from $^{14}$C-choline-chloride (American Radiolabeled Chemicals, Inc. [ARC 0208, 55 mCi/mmol]) as described *(25).* $^{14}$C-arginine was purchased from Moravek, Inc. [MC-137, 338 mCi/mmol], $^{14}$C-citrulline from American Radiolabeled Chemicals, Inc. [ARC 0508, 55 mCi/mmol] and $^{3}$H-ornithine hydrochloride from PerkinElmer Health Sciences, Inc. [NET1212, 21.4 Ci/mmol].

#### Construction of expression strains

The *arcX* genes were PCR-amplified from the genome of *Lactococcus lactis* IL1403 with primers *arcX*-Fw and *arcX*-rev (see Table S1), using Phusion HF DNA polymerase (Thermo Fisher Scientific, Inc.). The *arcA* and *arcB* PCR inserts, and the pNZcLIC*oppA* vector (*37*), were digested with *Nco*I and *Bam*HI and subsequently ligated to yield pNZ*arcA* and pNZ*arcB.* These vectors contain the corresponding genes under the control of the nisin-inducible $P_{\mathrm{NIS}}$ promoter (*38*) and the genes have a cleavable 6His-tag at the N-terminus.

The *arcC1* and *arcD2* PCR inserts were used for ligation-independent cloning as described in (*39*). This yielded pNZ*arcC1* and pNZ*arcD2* with the genes under the control of the $P_{\mathrm{NIS}}$ promoter and with a cleavable 10His-tag at the N- and C-terminus, respectively. To construct the cysteine-less variant of *arcD2* (*arcD2*ΔC), two mutations were made, namely C395T and C487T. The *arcD2* gene was PCR-amplified from pNZ*arcD2* with uracil-containing primers (*arcD2*ΔC-X), using PfuX7 DNA polymerase (40). The two amplified fragments were ligated with USER enzyme (New England Biolabs, Inc.) to create the pNZ*arcD2*ΔC vector.

The pNZ*arcA*, pNZ*arcB* and pNZ*arcC1* vectors were transformed into *L. lactis* NZ9000 (*38*), while pNZ*arcD2* and pNZ*arcD2*ΔC were transformed into *L. lactis* JP9000 Δ*arcD1D2* (*35*) (Table S2). The pRsetB-PercevalHR plasmid was a gift from professor Gary Yellen (Addgene plasmid #49081, *25*) and transformed into *E. coli* BL21-DE3 (Table S2).

**Table S1. Primers used for cloning**

| Primer | Sequence (5' -> 3') |
|---|---|
| *arcA*-Fw | AAACCATGGGTATGAACAATGGAATTAAATGTTAACTCAGAAATTGGG |
| *arcA*-Rv | ATAAGGATCCCAAATCTTCACGCCAAAGTGGTTGTGAC |
| *arcB*-Fw | AAACCATGGGTATGACATCACCACTTATTACAAAAGCAGAAGTAAAC |
| *arcB*-Rv | ATAAGGATCCTTTAAAATCTTCAGGAACTGCTGGGATAAATAAATTAC |
| *arcC1*-Fw | ATGGTGAGAATTTATATTTTCAAGGTGTAAAACGTATTGTTGTAGCCC |
| *arcC1*-Rv | TGGGAGGGTGGGATTTTCATTAAGGAACAATTCTTGTTCCACTAC |
| *arcD2*-Fw | ATGGGTGGTGGATTTGCTATGGAAAACAAGAAAACAAAAGGG |
| *arcD2*-Rv | TTGGAAGTATAAATTTTCGTAGCCTAGTAATCCCCG |
| *arcD2*ΔC-1 | AGCTGTAATUATGATTACTTATGCTTTGGTTGGAGC |

| | |
|---|---|
| *arcD2*ΔC-2<br>*arcD2*ΔC-3<br>*arcD2*ΔC-4 | AGACCAGAAUTGCTAAAACTCCTAGTAAGG<br>ATTCTGGTCUTGATTACTGGCGCAAAATCAGGAAC<br>AATTACAGCUGTTGCCATATAAATAAAGAC |

**Table S2. Strains used in this study**

| Strain | Genotype | Vector | Reference |
|---|---|---|---|
| *L. lactis* NZ9000<br>*L. lactis* NZ9000-A<br>*L. lactis* NZ9000-B<br>*L. lactis* NZ9000-C1 | MG1363 with *nisRK* in *pepN* locus | <br>pNZ*arcA*<br>pNZ*arcB*<br>pNZ*arcC1* | (*38*)<br>This study<br>This study<br>This study |
| *L. lactis* JP9000<br>*L. lactis* JP9000-D2<br>*L. lactis* JP9000-2ΔC | MG1363 with *nisRK* in pseudo_10 locus | <br>pNZ*arcD2*<br>pNZ*arcD2*ΔC | (*35*)<br>This study<br>This study |
| *E. coli* BL21(DE3) | $F^-$ *ompT gal dcm lon* $hsdS_B(r_B^- m_B^-)$ λ(DE3 [*lacI lacUV5-T7 ind1 sam7 nin5*]) | pRsetB-PercevalHR | This study |
| *L. lactis* OPU401 | NZ9000 Δ*opuA* | pNZ*opuA*His | (*33*) |

**Expression of genes and preparation of cell lysates and membrane vesicles**

*L. lactis* cells were grown in rich medium [2 % (w/v) Gistex from Brenntag AG], 65 mM NaPi, pH 7.0, 1 % (w/v) glucose) with 5 µg/mL chloramphenicol at 30 °C with stirring (200 rpm). The strains for ArcA, ArcB and ArcC1 production were grown as 3 L cultures in 5 L flasks and induced at an $OD_{600}$ of 0.5 with 0.05 % (v/v) of culture supernatant from a nisin A-producing strain (*38*). In contrast, the strains for ArcD2 and OpuA were grown as 2 L cultures in a 3 L fermentor stirred at 200 rpm with pH control (kept above pH 6.5 with 4 M KOH) and induced at an $OD_{600}$ of 2.0 with 0.05 % (v/v) of culture supernatant from a nisin A-producing strain. After induction, all strains were grown for an additional 2 hours before harvesting. Harvesting and washing was done by centrifugation (15 min, 6.000 x g, 4 °C) and resuspension of the cells in ice-cold 100 mM KPi, pH 7.0 (buffer A, Table S3). Finally, cells were centrifuged again and resuspended to an $OD_{600}$ of 100 in ice-cold 50 mM KPi, pH 7.0 (buffer B), flash-frozen in liquid nitrogen in aliquots of 50 mL and stored at -80 °C.

The preparation of cell lysate and membrane vesicles was done as follows. After cells were thawed on ice, 100 µg/mL DNAse and 2 mM $MgSO_4$ were added. Cells were lysed by high-pressure disruption (Constant Systems, Ltd.) with two passages at 39 kpsi and 4 °C. After lysis, 5 mM $Na_2$-EDTA (pH 8.0) and 1 mM PMSF (100 mM stock in isopropanol) were added and cell debris was removed by centrifugation (15 min, 22.000 x g, 4 °C). Next, the supernatant was centrifuged for 90 min at 125.000 x g at 4 °C. For ArcA, ArcB and ArcC1, the supernatant (containing cell lysate) was flash-frozen in liquid nitrogen in 10 mL aliquots and stored at -80 °C. For ArcD2 and OpuA, the pellets (containing membrane vesicles) were resuspended in ice-cold buffer B [with 20 % (v/v) glycerol for OpuA], to a 10 mg/mL protein concentration before flash-freezing (2 mL aliquots) and storage.

*E. coli* BL21-DE3 pRsetB-PercHR cells were grown in lysogeny broth (LB) with 100 µg/mL ampicillin. 0.5 L cultures were grown in 5 L flasks at 30 °C with shaking (200 rpm) to an $OD_{600}$ of 0.7, after which they were cooled to 22 °C, induced with 100 µM isopropyl β-D-1-thiogalactopyranoside (IPTG) and grown for an additional 72 hours before harvesting as described (*25*). Harvesting and washing was done by centrifugation (15 min, 6.000 x g, 4 °C) and resuspension in ice-cold 50 mM NaPi, pH 7.0 with 100 mM NaCl. Finally, cells were centrifuged again and resuspended to an $OD_{600}$ of 100 in ice-cold 20 mM NaPi, pH 7.0 with 100 mM NaCl, flash-frozen in liquid nitrogen in 50 mL aliquots and stored at -80 °C. Cell lysate was prepared by adding 250 µg of DNAse to thawed cells, after which they were lysed by high-pressure disruption in a single passage at 25 kpsi and 4 °C. After lysis, 0.1 mM PMSF was added and cell debris was removed by centrifugation for 60 min at 145.000 x g at 4 °C. The supernatant (containing cell lysate) was flash-frozen in liquid nitrogen in 15 mL aliquots and stored at -80 °C.

**Table S3. Buffers used in this study**

| Buffer | Composition |
|---|---|
| A | 100 mM KPi, pH 7.0 |
| B | 50 mM KPi, pH 7.0 |
| C | 50 mM KPi, pH 7.0 with 200 mM NaCl |
| D | 50 mM KPi, pH 7.0 with 100 mM NaCl |
| E | 25 mM KPi, pH 8.0 with 500 mM NaCl plus 5 % (v/v) glycerol |
| F | 50 mM KPi, pH 7.0 with 200 mM KCl |
| G | 100 mM KPi, pH 7.0 with 0.5 mM ornithine |
| H | 50 mM NaPi, pH 7.0 |
| I | 100 mM KPi, pH 7.0 with 250 mM KCl |

**Purification of ArcA, ArcB and ArcC1**

All protein purification and handling steps were performed on ice or at 4 °C, except when specified otherwise. $Ni^{2+}$-Sepharose resin was pre-equilibrated in 50 mM KPi, pH 7.0, with 200 mM NaCl (buffer C) with either 10 mM imidazole for ArcA and ArcB or 5 mM imidazole and 10 % (v/v) glycerol for ArcC1. Cell lysate was thawed on ice, added to the $Ni^{2+}$-Sepharose resin (0.5 mL bed volume per 10 mg total protein) and nutated for 1 hour. The mixture was poured over a polyprep column (Bio-Rad Laboratories, Inc.), after which the resin was washed with 20 column volumes of buffer C with 50 mM imidazole [plus 10 % (v/v) glycerol for ArcC1]. Proteins were then eluted with 3 column volumes of buffer C with 500 mM imidazole [plus 10 % (v/v) glycerol for ArcC1]. The most concentrated fractions were run on a Superdex 200 Increase 10/300 GL size-exclusion column (GE Healthcare) in 50 mM KPi, pH 7.0 with 100 mM NaCl (buffer D) [plus 10 % (v/v) glycerol for ArcC1]. Protein containing fractions were pooled and concentrated to 4-5 mg/mL in a Vivaspin 500 (30.000 kDa) centrifugal concentrator (Sartorius AG), after which they were aliquoted, flash-frozen in 100 µL aliquots and stored at -80 °C.

Additionally, ArcA, ArcB and ArcC1 have been purified in 50 mM NaPi, pH 7.0 instead of 50 mM KPi pH 7.0 to allow reconstitutions devoid of potassium ions (*Protocol A6*, see below); the system is also fully functional in sodium ion-based buffers.

**Purification of PercevalHR**

PercevalHR was purified in a manner similar to ArcA, ArcB and ArcC1, except that different buffers were used. $Ni^{2+}$-Sepharose resin was pre-equilibrated in 25 mM KPi buffer, pH 8.0 with 500 mM NaCl with 5 % (v/v) glycerol (buffer E) plus 10 mM imidazole. The resin was washed with buffer E plus 25 mM imidazole and protein was eluted with buffer E plus 250 mM imidazole. The most concentrated fractions were run on a Superdex 200 Increase 10/300 GL size-exclusion column (GE Healthcare) in 10 mM NaPi, pH 7.4 with 150 mM NaCl and 5 % (v/v) glycerol. Protein containing fractions (1-2 mg/mL) were aliquoted in volumes of 50 µL, flash-frozen and stored at -80 °C.

**Enzymatic assays for ArcA and ArcB**

Activity of ArcA and ArcB was measured with the COLDER assay (*41*). In brief, either 2 µg/mL ArcA, or 0.25 µg/mL ArcB was incubated in buffer B at 30 °C for 3 min, in a total volume of 275 µL. To start the reaction, varying concentrations of either L-arginine for ArcA (0 to 480 µM L-arginine), or L-ornithine plus carbamoyl-phosphate for ArcB (0 to 10 mM L-ornithine plus 0 to 10 mM carbamoyl-phosphate) were added. 200 µL of COLDER solution (20 mM 2,3-butanedione monoxime, 0.5 mM thiosemicarbazide, 2.25 M phosphoric acid, 4.5 M sulfuric acid and 1.5 mM ammonium iron(III) sulfate) was pipetted into each well of a 96-well flat-bottom transparent polystyrene plate (Greiner Bio-One International GmbH), to which 50 µL of reaction mixture was added at given time intervals to stop the enzymatic conversion. Additionally, a set of calibration samples with citrulline concentrations from 0 to 250 µM was added into the wells in the plate. To allow color development, the plate was sealed with thermo resistant tape (Nalge Nunc International) and incubated at 80 °C for 20 min in a block heater (Stuart). Afterwards, the plate was cooled down to room temperature for 30 min, the condensate was centrifuged (1 min, 1000 x g, 20 °C) and the absorbance of the solutions in the wells was measured at 540 nm in a platereader (BioTek Instruments, Inc.). Enzyme activity (in nmol citrulline x $min^{-1}$ x mg $protein^{-1}$) was determined by the formula:

$$Act_{enz} = \frac{\Delta enz}{\Delta cal} * \frac{1}{C_{enz} * Vol_{rmx}} \quad [1]$$

where Δenz and Δcal are the slopes of the enzyme and calibration curves, respectively, in AU/min and AU/nmol citrulline; $C_{enz}$ is the final concentration of enzyme in mg/mL and $Vol_{rmx}$ is the volume of the reaction mixture in mL.

Stability measurements of ArcA and ArcB were performed as described above, with minor adjustments. In brief, ArcA and ArcB were diluted in either buffer B or 300 mM KPi, pH 7.0, with and without 200 mM glycine betaine, and incubated at 30 °C for 0, 1, 3 and 5 hours. To start the reaction, either 150 µM arginine (for ArcA) or 5 mM carbamoyl-phosphate plus 5 mM citrulline (for ArcB) were added.

**Enzymatic assays for ArcC1**

The activity of ArcC1 was measured from changes in ATP/ADP ratio with PercevalHR. 3.3 µg/mL ArcC1 was incubated in buffer B supplemented with 5 mM of $MgSO_4$ and 10 µg/mL of purified PercevalHR in 105.250-QS cuvettes (Hellma Analytics) in a FP-8300 spectrofluorometer (Jasco, Inc.). ADP was added in varying concentrations (0.1 to 10 mM) and the mixture was incubated at 30 °C for 5 minutes. To start the reaction, carbamoyl phosphate was added in varying concentrations (0.2 to 10 mM). The fluorescence spectrum of PercevalHR was measured by excitation from 400 ± 5 nm to 510 ± 5 nm, while the emission was recorded at 550 ± 5 nm. As the pH of the reaction mixture changes in time and PercevalHR is sensitive to pH, it was necessary to measure the pH changes and correct the Perceval HR readout accordingly. In the pH experiments PercevalHR was substituted with 0.1 µM pyranine and the fluorescence spectrum of pyranine was measured by excitation from 380 ± 5 nm to 480 ± 5 nm, while the emission was recorded at 512 ± 5 nm. Pyranine was calibrated as described under *Internal pH measurements*.

The PercevalHR signal was calibrated at a sensor concentration of 10 µg/mL in buffer B with varying pH values (from 6.6 to 7.6), containing a mixture of ATP and ADP at a total concentration of 5 mM and a total $MgSO_4$ concentration of 5 mM. The ATP/ADP ratio was plotted against the ratio of the two excitation peaks at 430 nm and 500 nm and were fitted using the Hill equation:

$$\frac{F500nm}{F430nm} = start + (end - start)\frac{(\frac{ATP}{ADP})^n}{k^n + (\frac{ATP}{ADP})^n} \qquad [2]$$

where *k* is the apparent affinity constant; *n* is the Hill coefficient; *start* and *end* refer to the *y*-values at the vertical asymptotes. The Hill coefficient was constrained to 1. When the parameters of datasets recorded at varying pH values were compared, it was evident that only *start* and *end* were affected by pH, *k* remained constant. The *start* and *end* values were plotted against the pH and were fitted by using the following logistic equations (Fig. S2C):

$$start = y_0 + A * e^{R_0 * pH}, \qquad end = y_0 + A * e^{R_0 * pH} \qquad [3]$$

Where $y_0$, A and $R_0$ are the fit parameters. The resulting equations were incorporated into Eq. 2 to yield a formula in which the ATP/ADP ratio is dependent on the pH of the solution and the ratio of the excitation peaks at 430 nm and 500 nm of PercevalHR:

$$\frac{ATP}{ADP} = \frac{4.11 * \left(\frac{F500nm}{F430nm} - \left(-0.021 + 2 * 10^{-6} * e^{1.76 * pH}\right)\right)}{\left(-1.15 + 3.5 * 10^{-3} * e^{0.96 * pH}\right) - \frac{F500nm}{F430nm}} \qquad [4]$$

Stability measurements of ArcC1 were performed as described above, with minor adjustments. In brief, ArcC1 was diluted in buffer B or 300 mM KPi, pH 7.0, with and without 200 mM glycine betaine, and incubated at 30 °C for 0, 1, 3 and 5 hours. After incubation, 5 mM of ADP, 5 mM of $MgSO_4$ and either PercevalHR or pyranine were added. After 5 minutes of incubation at 30 °C, the reaction was started with the addition of 5 mM of carbamoyl phosphate. For the experiments performed in 300 mM KPi pH 7.0 a new calibration of both PercevalHR and pyranine was performed, resulting in the following equation:

$$\frac{ATP}{ADP} = \frac{6.59 * \left(\frac{F500nm}{F430nm} - \left(-0.214 + 8.2 * 10^{-5} * e^{1.29 * pH}\right)\right)}{\left(-0.52 + 6.3 * 10^{-4} * e^{1.15 * pH}\right) - \frac{F500nm}{F430nm}} \qquad [5]$$

**Purification of ArcD2 and OpuA**

Membrane vesicles were quickly thawed and diluted to a total protein concentration of 3 mg/mL for OpuA and 7 mg/mL for ArcD2 in 50 mM KPi, pH 7.0 plus 200 mM KCl (buffer F) containing 20 % (v/v) glycerol in case of OpuA. 0.5 % (w/v) *n*-dodecyl-β-D-maltoside (DDM) was added to the vesicles for solubilization and the mixture was nutated for either 30 min (ArcD2) or 60 min (OpuA). Unsolubilized material was removed by centrifugation (20 min, 270.000 x g, 4 °C). $Ni^{2+}$-Sepharose resin (0.5 mL of $Ni^{2+}$-Sepharose resin per 20 mg total protein for OpuA or 0.25 mL of $Ni^{2+}$-Sepharose resin per 10 mg total protein for ArcD2) was pre-equilibrated in buffer F with 10 mM imidazole plus 0.03 % (w/v) DDM. The supernatant was diluted 1.6x fold (ArcD2) or 2.5x fold (OpuA) to reduce the DDM concentration and then added to the $Ni^{2+}$-Sepharose column material and nutated for 1 hour at 4 °C. The mixture was poured into a poly-prep column, after which the resin was washed with 20 column volumes of buffer F containing 50 mM imidazole plus 0.02 % (w/v) DDM and 20 % (v/v) glycerol in case of OpuA. Proteins were eluted in 3 column volumes of buffer F with 500 mM imidazole plus 0.02 % (w/v) DDM and 20 % (v/v) glycerol in case of OpuA.

**Light scattering for oligomeric state determination**

$Ni^{2+}$-Sepharose/size-exclusion chromatography-purified fractions of ArcA, ArcB, ArcC and ArcD2 were analyzed on a second Superdex 200 Increase 10/300 GL size-exclusion column (GE Healthcare) in buffer D [with 0.02 % (w/v) DDM for ArcD2], which was coupled to a multi-angle light scattering system with detectors for absorbance at 280 nm (Agilent Technologies, Inc.), static light scattering (Wyatt Technology Corporation) and differential refractive index (Wyatt Technology Corporation). Data analysis was performed with the ASTRA software package (Wyatt Technology Corporation), using a value for the refractive index increment $(dn/dc)_{protein}$ of 0.180 mL/mg and $(dn/dc)_{detergent}$ of 0.143 mL/mg (*42*).

**Co-reconstitution of ArcD2 and OpuA**

Synthetic lipids were mixed from chloroform stocks in the ratio of either 50 mole% DOPE, 12 mole% DOPC and 38 mole% DOPG or 50 mole% DOPE, 37 mole% DOPC and 13 mole% DOPG. Lipids were dried in a rotary vacuum setup (Büchi Labortechnik AG), dissolved in diethyl ether, dried again and rehydrated in buffer B to a final lipid concentration of 20 mg/mL. Dissolved lipids, cooled with ice water, were sonicated with a tip sonicator (Sonics and Materials, Inc.) (15 sec on, 45 sec off, 70 % amplitude, 16 cycles), frozen-thawed 3 times, alternating between liquid nitrogen and (a water bath at) room temperature, and extruded 13 times through a 400 nm pore size polycarbonate filter (Avestin Europe GmbH) to obtain liposomes. Using an established protocol (*34*), ArcD2 and OpuA were co-reconstituted in preformed liposomes at a protein to lipid ratio of 1:2:400 (w/w), respectively. The liposomes were first diluted 5 times to a final concentration of 4 mg/mL in buffer B with 25% (v/v) glycerol [final concentration 20 % (v/v)] and then destabilized by a stepwise titration with 10 % (v/v) Triton X-100, until the membrane was saturated with detergent ($R_{sat}$; *43*), after which the membrane proteins were added. The purified proteins and destabilized liposomes were mixed for 15 min at 4 °C, after which detergent was removed by adding SM2 biobeads (600 mg per 20 mg of lipids) in three equal aliquots with 15 min incubation in between each addition. After the third addition, the mixture was incubated overnight, which was followed by one additional addition (200 mg per 20 mg of lipids) of SM2 biobeads and incubation for one hour. Finally, the proteoliposomes were collected by centrifugation (2 hours for 38% (w/w) DOPG lipids or 4 hours for 13% (w/w) DOPG lipids, 125.000 x g, 4 °C) and resuspended in 200 µL per 20 mg of lipids, yielding a final lipid concentration of 100 mg/mL. For the $^{14}C$-arginine transport assay (see transport assays), reconstitution was done similarly as above, except that ArcD2 was reconstituted at a protein to lipid ratio of 2:400 (w/w). Additionally, the proteoliposomes were diluted in buffer B without glycerol and centrifugation was done for 30 min, 325.000 x g, 4 °C.

**Encapsulation of the arginine breakdown pathway**

*Protocol A1.* The ArcD2- and OpuA-containing vesicles used for most studies were composed of 50 mole% DOPE, 12 mole% DOPC plus 38 mole% DOPG. The proteoliposomes (66 µL, 6.6 mg of lipid) containing ArcD2 and OpuA were mixed in buffer B with 1 µM ArcA, 2 µM ArcB, 5 µM ArcC1, 5 mM ADP, 5 mM $MgSO_4$, 0.5 mM ornithine and optionally 1.6 - 2.9 µM PercevalHR or 100 µM pyranine in a total volume of 200 µL; the final liposome concentration is 33 mg of lipid/mL. This yields a final buffer of 50 mM KPi pH 7.0 plus 25 mM NaCl (carried over with the purified ArcA, ArcB and ArcC1) or 40 mM NaCl when PercevalHR is also included (see Table S4). The enzymes, metabolites and dyes were encapsulated by five freeze-thaw cycles in a 0.5 ml Eppendorf tube, alternating between liquid nitrogen and a 10 °C water bath, with vortexing of the tube before freezing. Next, the vesicles were extruded 13 times through a 400 nm pore size polycarbonate filter; the extruder was pre-washed in buffer A with 0.5 mM ornithine (=buffer G). This procedure homogenizes the vesicles further and makes it likely that necessary components are present in all layers and compartments. The vesicles with encapsulated pyranine were then separated from free pyranine by running them over a 22 cm long Sephadex G-75 (Sigma) column in buffer G at 4 °C. To remove the residual external compounds, the vesicles were diluted to 6 mL in buffer G, collected by centrifugation (20 min, 325.000 x g, 4 °C) and washed with buffer G (6 mL), after which the vesicles were centrifuged and resuspended in 40 µL per 6.6 mg of lipid, yielding a final concentration of 165 mg of lipid/mL. Vesicles were kept on ice before subsequent measurements. Importantly, the ratio of the components inside the vesicles is very similar to the ratio in solution prior to encapsulation (Fig. S6).

*Protocol A2.* Like *Protocol 1* except that the proteoliposomes were mixed with internal components in 60 mM KCl, yielding a final buffer of 15 mM KPi pH 7.0 plus 25 mM NaCl and 40 mM KCl.

*Protocol A3.* Like *Protocol 1* except that the vesicles were loaded with 15 mM ADP plus equimolar concentrations of $MgSO_4$.

*Protocol A4.* Like *Protocol 1* except that the vesicles were extruded through a 200 nm pore size polycarbonate filter.

*Protocol A5.* Like *Protocol* 1 except that the vesicles were loaded with 10 mM ATP, 10 mM $MgSO_4$, 24 mM creatine phosphate and 2.4 mg/mL creatine kinase in buffer B (*34*).

*Protocol A6.* Like *Protocol 1* except that the proteoliposomes containing ArcD2 and OpuA were first diluted in 6 ml of 50 mM NaPi, pH 7.0 (buffer H), collected by centrifugation (20 min, 325.000 x g, 4 °C) and resuspended in 66 µl of buffer H. They were then mixed in buffer H with the components mentioned above and the ArcA, ArcB and ArcC1 purified in 50 mM NaPi instead of 50 mM KPi. Additionally, the extruder was pre-washed and vesicles were resuspended in buffer H with L-ornithine. This encapsulation yields only sodium and no potassium ions inside the vesicles.

**Table S4. Average number of molecules per vesicle**

| Compound | Internal concentration | Molecules per vesicle (84 nm radius) | Molecules per vesicle (225 nm radius) |
|---|---|---|---|
| Ornithine | 0.5 mM | 740 | 14,500 |
| ADP | 5 mM | 7,400 | 144,600 |
| $Mg^{2+}$ | 5 mM | 7,400 | 144,600 |
| Pyranine | 100 µM | 150 | 2,900 |
| PercevalHR | 2.9 µM | 4 | 83 |
| Phosphate | 50 mM | 73,700 | 1,446,200 |
| $K^+$ | 50 mM | 73,700 | 1,446,200 |
| $Na^+$ | 25 / 40 mM | 36,800 / 59,000 | 723,100 / 1,157,000 |
| $Cl^-$ | 25 / 40 mM | 36,800 / 59,000 | 723,100 / 1,157,000 |
| ATP/ADP ratio of 3 (without GB uptake) | 3.75 / 1.25 mM | 5,500 / 1,800 | 108,500 / 36,200 |
| ATP/ADP ratio of 2 (with GB uptake) | 3.33 / 1.67 mM | 4,900 / 2,500 | 96,400 / 48,200 |
| Glycine betaine (imported) | 50 mM | 73,700 | 1,446,200 |

**Cryo-EM analysis of vesicles**

The vesicles with encapsulated enzymes, metabolites (and sensors) were vitrified and imaging was done on a FEI Tecnai T20, 200 keV; Cryo-stage Gatan model 626. Samples were prepared under isosmotic conditions and images were recorded under low-dose conditions (*44*). Approximate diameters of the vesicles were measured in ImageJ. The diameters were converted to internal volume by assuming spherical vesicles, multiplication by abundance and re-normalization.

**Transport assays**

*Protocol B1.* The vesicles with encapsulated enzymes and metabolites were diluted to a final concentration of 1.67 mg/mL in buffer A with 250 mM KCl (buffer I). Glycine betaine was added at a final concentration of 18 µM, of which 2 % (mol/mol) was $^{14}C$-radiolabeled. The mixture was incubated for 30 minutes at 30 °C. The internal ATP production was then started by addition of 20 mM arginine and samples of 50 µL were taken at given time intervals. Samples were diluted in 2 mL of ice-cold buffer I and filtered over 0.45 µm pore size cellulose nitrate filters to stop the transport assay. The filter was then washed with another 2 mL of buffer I. Radioactivity on the filter was quantified by liquid scintillation counting using Ultima Gold MV scintillation fluid (PerkinElmer) and a Tri-Carb 2800TR scintillation counter (PerkinElmer). The pore size of the filters is larger than the diameter of the vesicles, but the filters retain more than 99% of the vesicles and allow for rapid filtration (*34*).

*Protocol B2*. The transport of $^{14}C$-arginine was measured similarly, except that proteoliposomes (66 µL, 6.6 mg of lipid) with only ArcD2 in the membrane, and L-ornithine or L-citrulline (100 μM, 1 mM or 10 mM) in buffer B in the vesicle lumen, were used (encapsulation was done by 5 freeze-thaw cycles in a total volume of 200 μL). The proteoliposomes were first extruded 13 times through a 400 nm pore size polycarbonate filter, then 13 times through a 200 nm filter and diluted to 6 mL in buffer B with or without the same concentration of L-ornithine or L-citrulline as on the inside. Proteoliposomes were collected by centrifugation (20 min, 225.000 x g, 4 °C) and either washed with buffer B (6 mL), centrifuged again and resuspended in 30 μL buffer B per 6.6 mg of lipid, or directly resuspended in buffer B with the appropriate concentration of L-ornithine or L-citrulline, yielding a final concentration of 220 mg of lipid/mL. For the transport assay, proteoliposomes were diluted to a final concentration of 2.2 mg of lipid/mL, in buffer B with 10 μM arginine, of which 10 % (mol/mol) was $^{14}C$-radiolabeled, and 100 μL samples were taken at given time intervals. To impose a membrane potential, proteoliposomes in buffer B, were diluted 100-fold in 50 mM NaPi ($\Delta\Psi$ = -120 mV), pH 7.0; 48.15 mM NaPi plus 1.85 mM KPi, pH 7.0 ($\Delta\Psi$ = -80 mV); or 39.6 mM NaPi plus 10.4 mM KPi, pH 7.0 ($\Delta\Psi$ = -40 mV), each supplemented with 1 μM of the potassium ionophore valinomycin.

**ATP:ADP ratio measurements with PercevalHR inside the vesicles**

*Calibration.* Purified PercevalHR and nucleotides (ATP and ADP) were encapsulated in liposomes. The encapsulation mixture contained liposomes at 7.5 mg of lipids/mL, 1.6 - 2.9 μM PercevalHR, 5 mM of nucleotides in varying ratios and 5 mM $MgSO_4$ in buffer B. The samples were frozen-thawed 5 times, extruded 13 times through a 400 nm pore size polycarbonate filter, centrifuged twice (20 min, 225.000 x g, 4 °C) and finally resuspended in buffer B to a concentration of 167 mg/mL. The liposomes were diluted in buffer I with 10 μM carbonyl cyanide-4-(trifluoromethoxy) phenylhydrazone (FCCP) to a final concentration of 3.34 mg lipids/mL in 105.250-QS cuvettes (Hellma Analytics) in a FP-8300 spectrofluorometer (Jasco, Inc.) and incubated for 3 min at 30 °C. The fluorescence spectrum of PercevalHR was measured by excitation from 400 ± 5 nm to 510 ± 5 nm, while the emission was recorded at 550 ± 5 nm. The encapsulated ATP/ADP ratio was plotted against the ratio of the peaks at 500 nm and 430 nm. Equation 2 was re-written with n = 1, to obtain the following equation:

$$\frac{ATP}{ADP} = \frac{k * (\frac{F500nm}{F430nm} - start)}{end - \frac{F500nm}{F430nm}} \qquad [6]$$

By fitting Eq. 6 to the data points (Fig. S2D) the following values were obtained: *k* = 3.02, start = 0.46, end = 1.30.

*Protocol B3.* The vesicles with encapsulated enzymes, metabolites and PercevalHR were diluted in buffer I with 10 μM FCCP to a final concentration of 3.34 mg of lipid/mL in 105.250-QS cuvettes (Hellma Analytics) in a FP-8300 spectrofluorometer (Jasco, Inc.) and incubated for 30 min at 30 °C. To start ATP production, 5 mM arginine was added, and after 30 min of incubation glycine betaine (0, 180 μM or 3.6 mM) was added. The fluorescence spectrum of PercevalHR was measured continuously, as described above. The ATP/ADP ratio was obtained from the ratio of the peaks at 500 nm and 430 nm, using equation 3. Representative traces are shown in the figure panels, but replicate experiments have been performed multiple times (*n*); n values are given in the figure legends.

**Internal pH measurements with pyranine**

*Calibration.* 100 µM pyranine was encapsulated in liposomes at varying pH between 6.0 and 8.0 in 50 mM KPi. The samples were frozen-thawed 5 times, extruded 13 times through a 400 nm pore size polycarbonate filter and run over a 22 cm long Sephadex G-75 (Sigma) column. The vesicles were collected by centrifugation (20 min, 325.000 x g, 4 °C), washed once, centrifuged and resuspended to a concentration of 167 mg of lipid/mL. The liposomes were diluted in 50 mM KPi at varying pH to a final concentration of 3.34 mg lipids/mL in 105.250-QS cuvettes (Hellma Analytics) in a FP-8300 spectrofluorometer (Jasco, Inc.) and incubated for 3 min at 30 °C. The fluorescence spectrum of pyranine was measured by excitation from 380 ± 5 nm to 480 ± 5 nm, while the emission was recorded at 512 ± 5 nm. The fluorescence spectrum of 0.1 µM pyranine in solution at varying pH between 6.0 and 9.0 in 50 mM KPi was also measured. The data points from encapsulated pyranine in unshocked vesicles overlapped perfectly with the in solution data, therefore the pH in solution was plotted against the ratio of the peaks at 450 nm and 405 nm and fitted using the following logistic function:

$$y = \frac{L}{1+e^{-k*(x-x_0)}} \quad [7]$$

where $L$ is the curve's maximum value; $k$ is the logistic growth rate and $x_0$ is the $x$-value of the sigmoid's midpoint. Equation 7 was re-written with the following parameters: x = pH and y = F450nm / F405nm to obtain the following equation:

$$pH = \frac{\ln\left(L * \frac{F405nm}{F450nm} - 1\right)}{-k} + x_0 \quad [8]$$

By fitting Eq. 7 to the data points in 50 mM KPi (Fig. S4B), the following values were obtained: $L = 3.60$, $k = 2.38$ and $x_0 = 7.84$.

When vesicles are exposed to an osmotic upshift by the addition of 250 mM KCl, the internal KPi concentration increases to ~300 mM. The high salt concentration shifts the calibration of pyranine, and therefore additional calibration curves were made with pyranine in 300 mM KPi at pH values ranging from 6.0 and 9.0. This data was fitted to Eq. 7 (Fig. S4B), as above, to obtain the following values: $L = 3.70$, $k = 2.21$ and $x_0 = 7.57$. The calibration is also shifted when the vesicles were encapsulated with 15 mM Mg-ADP instead of 5 mM Mg-ADP (see *Protocol A3*). Therefore we also made calibration curves with pyranine in 300 mM KPi plus 15 mM Mg-ADP at pH values ranging from 6.0 to 7.5 (with 15 Mg-ADP and 300 mM KPi the salts precipitated above pH 7.5). This data was fitted to Eq. 7 (Fig. S4C) to obtain the following values: $L = 3.22$, $k = 2.16$ and $x_0 = 7.56$.

*Protocol B4.* The vesicles with encapsulated enzymes, metabolites and pyranine were diluted in buffer I to a final concentration of 3.34 mg lipids/mL in 105.250-QS cuvettes (Hellma Analytics) in a FP-8300 spectrofluorometer (Jasco, Inc.) and incubated for 30 min at 30 °C. To start the ATP production, 5 or 20 mM arginine was added and after 30 min of incubation, glycine betaine was added at a concentration of 0 or 180 µM. The fluorescence spectrum of pyranine was measured as indicated above, and the internal pH was obtained from Eq. 8. Representative traces are shown in the figure panels, but replicate experiments have been performed multiple times (*n*); n values are given in the figure legends.

**External pH measurements with pyranine**

*Calibration.* The fluorescence spectrum of pyranine was measured by excitation from 380 ± 5 nm to 480 ± 5 nm, while the emission was recorded at 512 ± 5 nm. The fluorescence spectrum of 0.1 µM pyranine in solution at varying pH between 6.0 and 8.5 in 10 mM KPi plus 355 mM KCl was measured. The pH in solution was plotted against the ratio of the peaks at 450 nm and 405 nm and fitted using equation 7 (Fig. S4D), from which the following values were obtained: $L$ = 3.27, $k$ = 2.23 and $x_0$ = 7.48.

*Protocol B5.* The vesicles with encapsulated enzymes and metabolites were diluted in 10 mM KPi pH 7.0 plus 355 mM KCl to a final concentration of 3.34 mg of lipids/mL in 105.250-QS cuvettes (Hellma Analytics) in a FP-8300 spectrofluorometer (Jasco, Inc.) and incubated for 30 min at 30 °C. To start the ATP production, 5 mM arginine was added. The fluorescence spectrum of pyranine was measured as indicated above, and the internal pH was obtained from Eq. 8. Representative traces are shown in the figure panels, but replicate experiments have been performed multiple times (*n*); n values are given in the figure legends.

**Amino acid and ammonia analysis**

*Protocol B6.* The vesicles with encapsulated enzymes and metabolites were diluted in buffer I, with or without glycine betaine, to a final concentration of 3.34 mg lipids/mL and incubated for 30 minutes at 30 °C. The assay was started by addition of 5 mM arginine and samples of 75 µL were taken at regular time intervals. Vesicles were removed from the samples by centrifugation (20 min, 225.000 x g, 4 °C) to stop the conversion of external amino acids. Subsequently, 50 µL of supernatant was mixed with 87.5 µL of 1 M boric acid, pH 9.0 (adjusted with 4 M KOH) and kept on ice before derivatization. 37.5 µL of 99.8% methanol and 1.5 µL of 42 mM diethyl ethoxymethylenemalonate (DEEMM) were added to the samples for derivatization, after which the samples were incubated for 30 min in a sonication bath at room temperature, followed by 2 hours of incubation in a heat block at 70 °C. The protocol for reverse-phase HPLC was adapted from (*45*). Amino acid samples were analyzed on a Shimadzu prominence HPLC system, containing a DGU-20A5R degassing unit, a LC-30AD solvent delivery unit, a SIL-30AC autosampler, a CTO-20AC column oven and an SPD-M20A UV-VIS/ Photodiode Array detector. A Shimadzu XR-ODS 3x75 mm C18 column was used to run the binary gradient with a flow rate of 0.9 mL/min and an injection volume of 5 µL. Eluent A was 25 mM acetate, pH 5.8, supplemented with 0.02 % (w/v) Na-azide. Eluent B was an 8:2 (v/v) mixture of acetonitrile and methanol. The gradient was as follows (all percentages are volume-based): start was at 94 % Eluent A and 6 % B; 87 % A and 13 % B at 2 min; 83 % A and 17 % B at 10 min; 71 % A and 29 %B at 11 min; 67 % A and 33 % B at 16 min; 40 % A and 60 % B at 16.1 min to 18 min; 94 % A and 6 % B at 18.1 min towards the end of the protocol at 20 min. The compounds were identified based on retention times and quantified using the external standard method.

**Membrane permeability by stopped-flow fluorescence-based measurements**

*Principle of the method.* To determine the permeability of solutes through the vesicle membrane, two independent and complementary fluorescence-based kinetic assays were used as described (*46*). The first assay reports volume changes of vesicles by means of calcein self-quenching fluorescence (*24*). The second assay monitors the pH variation in the vesicle lumen using the ratio-metric fluorophore pyranine (*47*). Both assays exploit the out-of-equilibrium relaxation kinetics of the vesicles after the increase of the external osmotic pressure (osmotic upshift), *i.e.*

the addition of a solute to the vesicle solution. After the osmotic upshift, the thermodynamic equilibrium is re-established by (i) water efflux (to re-equilibrate the chemical potential of water) and/or (ii) solute influx (to dissipate the solute concentration gradient) (*48*). The contribution of the two fluxes to the recovery kinetics depends on the relative permeability of water and the solute as described (*46*).

*Osmolyte solutions.* The 1 M stock solutions of Na-acetate, $NH_4Cl$ and $NH_4$-acetate were prepared by dissolving the salts into 100 mM KPi buffer; the pH was adjusted to 7.0 using 4 M NaOH. An empirical linear relation between concentration and osmolality for each solution was determined as described (*46*). The stock solutions were then diluted to an osmolality of ca. 300 mosmol/kg by mixing with the liposome solution (yielding 95 mM Na-acetate, 110 mM $NH_4Cl$ and 110 mM $NH_4$-acetate).

*Liposome preparation.* Liposomes were prepared as described under pH measurements, with minor adjustments. Calcein was added to the vesicle solution (2 mg of lipid in buffer A in a total volume of 1 mL) at a self-quenching concentration of 10 mM and enclosed by three cycles of freezing and thawing, alternating between liquid nitrogen and a 40°C water bath. Pyranine (300 µM pyranine, final concentration) was encapsulated similarly. Next, the liposomes were extruded 13 times through a 200 nm pore size polycarbonate filter and run over a 22 cm long Sephadex G-75 (Sigma) column in buffer A. Vesicles were collected and diluted to a total volume of 12 mL in buffer A. Empty liposomes for blank correction were prepared using the same procedure without addition of calcein or pyranine.

*Stopped-flow fluorescence measurements.* The permeability of the liposomes for KPi, KCl, Na-acetate, $NH_4Cl$ and $NH_4$-acetate was assessed by monitoring the quenching of the fluorescence of calcein and by determining the changes in the internal pH using pyranine as a probe. A stopped flow apparatus (SX20, Applied Photophysics Lim., Leatherhead, Surrey, UK) was used to measure fluorescence intensity kinetics upon imposition of an osmotic upshift to the liposomes filled with calcein or pyranine. The osmolyte (ca. 300 mosmol/kg after mixing) and the liposome solutions (pre-equilibrated with 250 mM KCl) were loaded each in two distinct syringes, forced through the mixer (1:1 mixing ratio and 2 ms dead time) and into the optical cell (20 µl volume and 2 mm path length). The temperature was set at 20°C using a water bath. The white light emitted by a Xenon arc lamp (150W) was passed through a high precision monochromator and directed to the optical cell via an optical fiber. The band pass of the monochromator was optimized and set to 0.5 nm (for calcein) or 1.4 nm (for pyranine) to prevent fluorophore photobleaching during the experiment. The fluorophores were excited at 495 nm (for calcein) or at both 405 nm and 453 nm (for pyranine). The emitted light, collected at 90°, was filtered by a Schott long-pass filter (cut-off wavelength at 515 nm) and detected by a photomultiplier tube (Hamamatsu R6095) with 10 µs time resolution. The voltage of the photomultiplier was automatically selected and kept constant during each set of experiments. The fluorescence intensity kinetics after the osmotic shock was recorded with logarithmically spaced time points to better resolve faster processes. For noise reduction, multiple acquisitions (3 for slow kinetics and 9 for fast kinetics) were performed for each experimental condition. Complete stopped- flow settings and acquired data processing are described in (*46*).

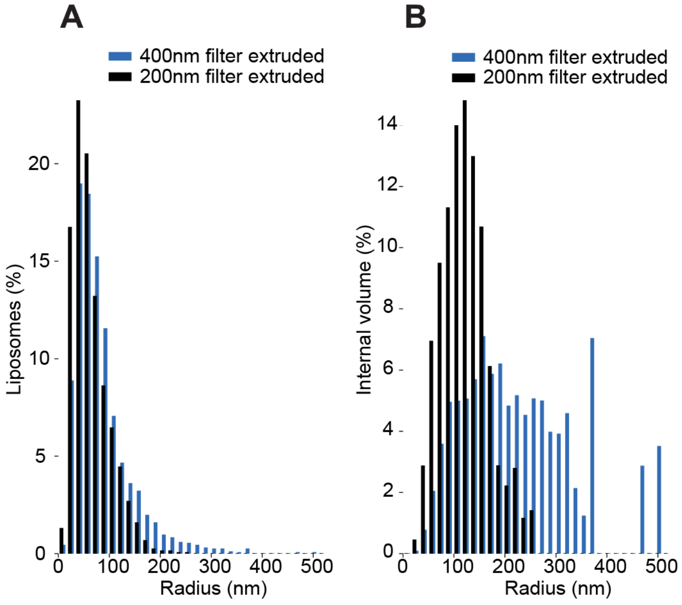

**Figure S1. Size distribution of the extruded lipid vesicles**
(A) Distribution of the radius of lipid vesicles extruded through a 400 nm (blue bars) and 200 nm (black bars) polycarbonate filter as estimated from CryoTEM micrographs. The diameter of 2090 vesicles (400 nm filter) and 2092 vesicles (200 nm filter) were measured using ImageJ. (B) Distribution of the internal volume, based on the distribution of radii, assuming that all vesicles are spherical. The distribution of the vesicles extruded through a 400 nm filter is wider and the fraction of multilamellar vesicles is higher compared to the 200 nm filter, but results obtained with both preparations were comparable (Fig. 3B and Fig. S5).

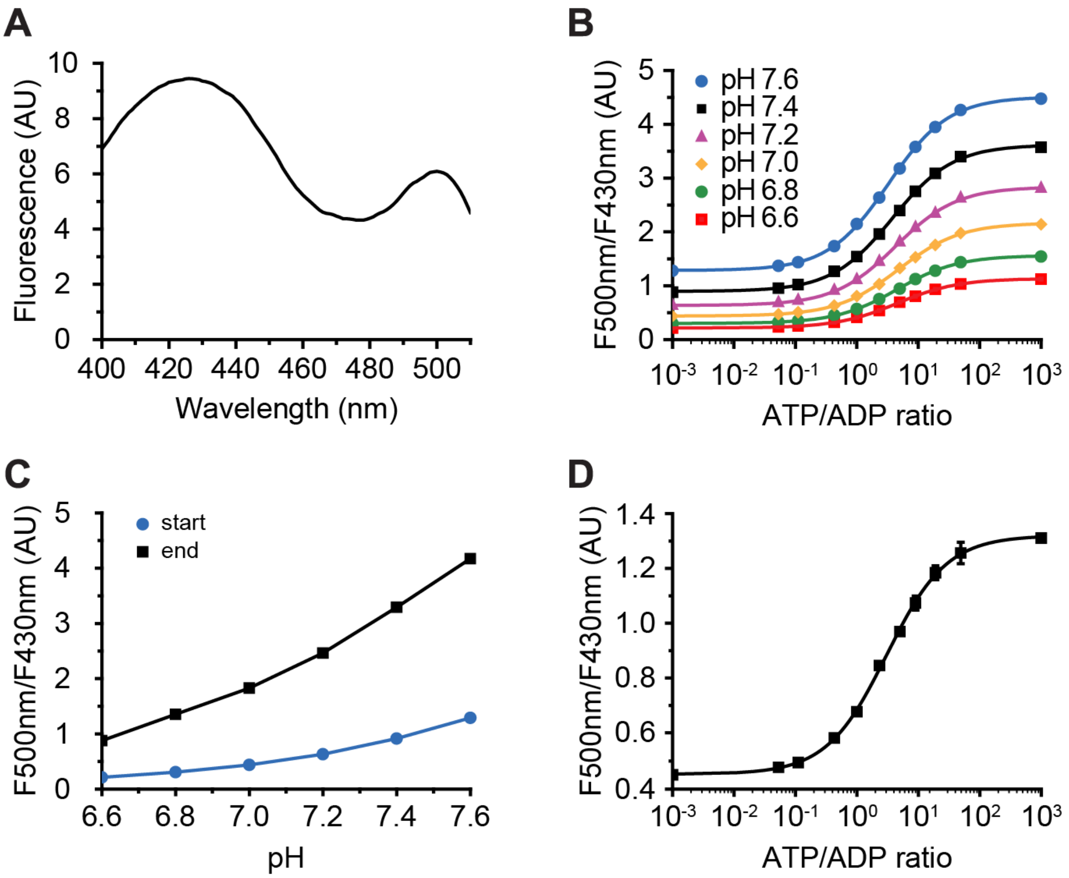

**Figure S2. Fluorescence-based ATP/ADP ratio sensor PercevalHR**

Fluorescence spectrum, calibration and pH dependency of PercevalHR measured in a FP-8300 spectrofluorimeter (Jasco, Inc.). (**A**) Excitation spectrum from 400 nm to 510 nm of PercevalHR encapsulated in the lipid vesicles with an emission wavelength of 550 nm, at equimolar ATP to ADP (2.5 mM each). The spectrum was corrected for background fluorescence. (**B**) Effect of pH on the readout of PercevalHR fluorescence. The ratio of the excitation peaks at 500 nm and 430 nm was measured for different ATP to ADP ratios, measured in 50 mM KPi pH 7.6 (blue circles), pH 7.4 (black squares), pH 7.2 (pink triangles), pH 7.0 (yellow diamonds), pH 6.8 (green circles), and pH 6.6 (red squares), each supplemented with 5.5 mM $MgSO_4$. (**C**) Plot of the *start* and *end* values against pH in 50 mM KPi, as fitted with Eq. 3. (**D**) Calibration of PercevalHR inside the lipid vesicles. The ratio of the excitation peaks at 500 nm and 430 nm changes when the ATP to ADP ratio is varied. Error bars indicate the standard deviation of two independent encapsulations. The data points were fit with the Hill equation (black line), as described in the materials and methods (n = 1, $k$ = 3.02, start = 0.46, end = 1.30).

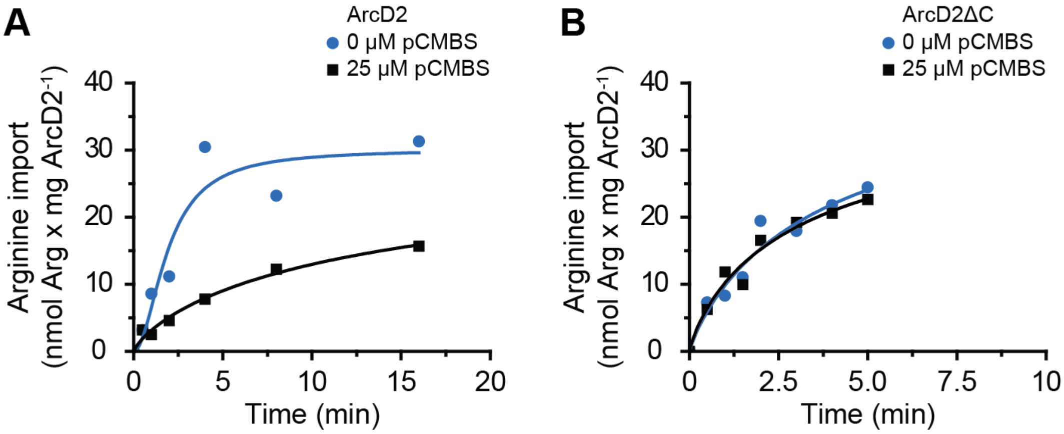

**Figure S3. pCMBS does not inhibit arginine/ornithine transport by ArcD2ΔC**
Radiolabeled arginine uptake by (**A**) wild-type ArcD2 and (**B**) cysteine-less ArcD2 (ArcD2ΔC) in the absence (blue circles) and presence (black squares) of 25 µM of pCMBS. The pCMBS was added 45 minutes prior to the start of the measurement to allow for the binding reaction to occur. The proteoliposomes were loaded with 0.5 mM L-ornithine and the final $^{14}$C-L-arginine concentration was 20 µM for the wild-type and 10 µM for the cysteine-less ArcD2. The presented data are obtained from a single experiment, but similar measurements (optimization of assay and labeling) were done multiple times.

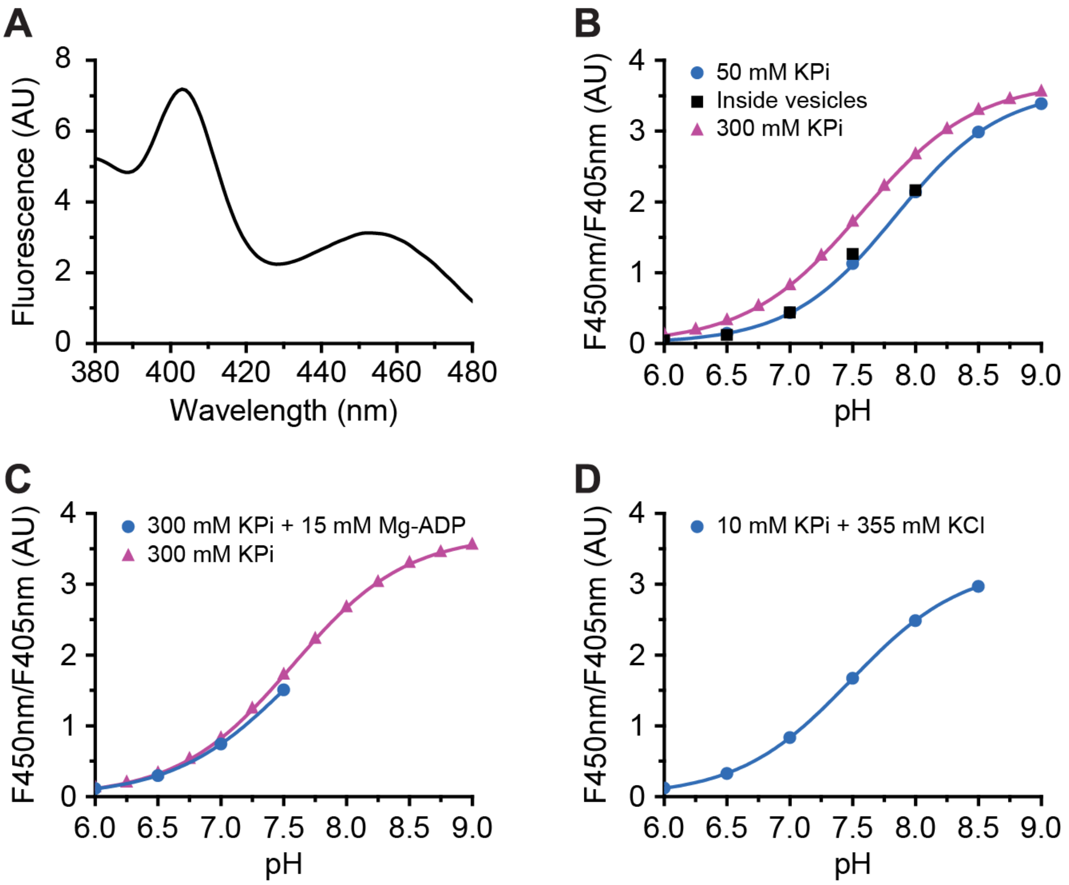

**Figure S4. Calibration of pyranine inside lipid vesicles**

(**A**) Excitation spectrum from 380 nm to 480 nm of pyranine inside the lipid vesicles with an emission wavelength of 512 nm, at pH 7.0. (**B**) Pyranine was measured at varying pH in 50 mM KPi (blue circles), inside the lipid vesicles containing 50 mM KPi (black squares) and in 300 mM KPi (pink triangles), using a FP-8300 spectrofluorimeter (Jasco, Inc.). The ratio of the excitation peaks at 450 nm and 405 nm changes as a function of pH. The data points in 50 and 300 mM KPi were fit with a logistic function (blue and pink traces), as described in the materials and methods section (50 mM KPi: $L = 3.60$, $k = 2.38$, $x_0 = 7.84$; 300 mM KPi: $L = 3.70$, $k = 2.21$, $x_0 = 7.57$). The data points inside the unshocked lipid vesicles perfectly match those in 50 mM KPi. (**C**) Similar to panel B, pyranine was measured at varying pH in 300 mM KPi with 15 mM Mg-ADP (blue circles) and 300 mM KPi (pink triangles). Data points in 300 mM KPi plus 15 mM Mg-ADP were fit (blue trace) to obtain: $L = 3.22$, $k = 2.16$ and $x_0 = 7.56$. (**D**) Similar to panel B, pyranine was measured at varying pH in 10 mM KPi plus 355 mM KCl (blue circles). Data points in 10 mM KPi plus 355 mM KCl were fit (blue trace) to obtain: $L = 3.27$, $k = 2.23$ and $x_0 = 7.48$.

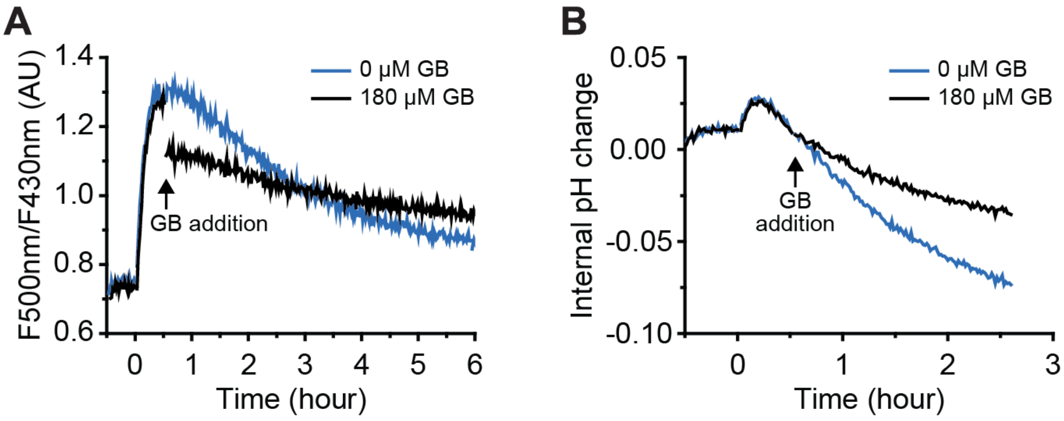

**Figure S5. ATP production and pH changes in arginine-metabolizing vesicles obtained by extrusion through 200 nm polycarbonate filters**

(**A**) Analogous to Fig. 3D, the effect of glycine betaine (GB) import on ATP production as measured by PercevalHR fluorescence (*Protocol B3*) in arginine-metabolizing vesicles exposed to an osmotic upshift (addition of 250 mM KCl) in the presence (black trace) and absence (blue trace) of 180 µM glycine betaine (added at t = 0.5 h); 5 mM arginine was added at t = 0 (*n*=2). (**B**) Analogous to Fig. 3G, the effect of glycine betaine on the internal pH measured by pyranine (*Protocol B4*) in arginine-metabolizing vesicles exposed to an osmotic upshift (250 mM KCl) in the presence (black trace) and absence (blue trace) of 180 µM glycine betaine (added at t = 0.5 h); 5 mM arginine was added at t = 0 (*n*=2).

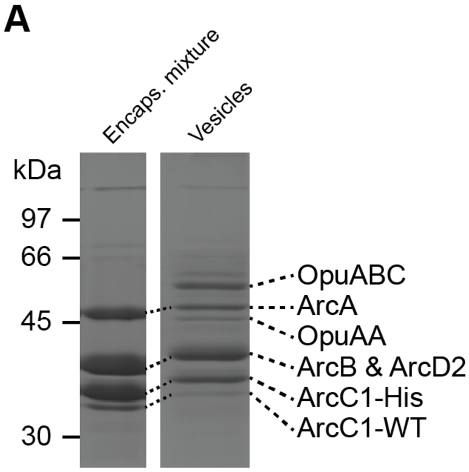

**B**

| | Encaps. mixture | Inside vesicles |
|---|---|---|
| ArcA | 1 | 1 |
| ArcB | 1.8 | 1.9 |
| ArcC1 | 1.6 | 1.4 |

**Figure S6. SDS-Polyacrylamide gel electrophoresis of purified and encapsulated proteins**

(**A**) On the left, encapsulation mixture containing 0.94 µg of ArcA, 2.4 µg of ArcB and 1.8 µg of ArcC1. On the right, the vesicles with encapsulated proteins and co-reconstituted ArcD2 and OpuA. OpuA consists of two separate proteins: OpuABC, containing the transmembrane domain fused to the substrate-binding domain; and OpuAA, the nucleotide-binding domain fused to a regulatory domain. Although ArcB and ArcD2 have a different monomeric molecular weight (40.9 and 56.7 kDa respectively), ArcD2 migrates at a similar position as ArcB, which has been observed for many other membrane proteins (*49*). The purification of his-tagged ArcC1 (ArcC1-His) always yields a small amount of wild type ArcC1 (ArcC1-WT), because the cell expresses ArcC1-WT at a basal level and heterooligomers are formed; the two proteins do migrate at a different position. (**B**) Quantification of the ratio of the ArcA, ArcB and ArcC1 before (encapsulation mixture) and after reconstitution (inside the vesicles) as analyzed with ImageJ (standard deviation from analyzing 4 independent loadings is 0.1). The ratio inside the vesicles does not differ significantly from that of the encapsulation mixture.

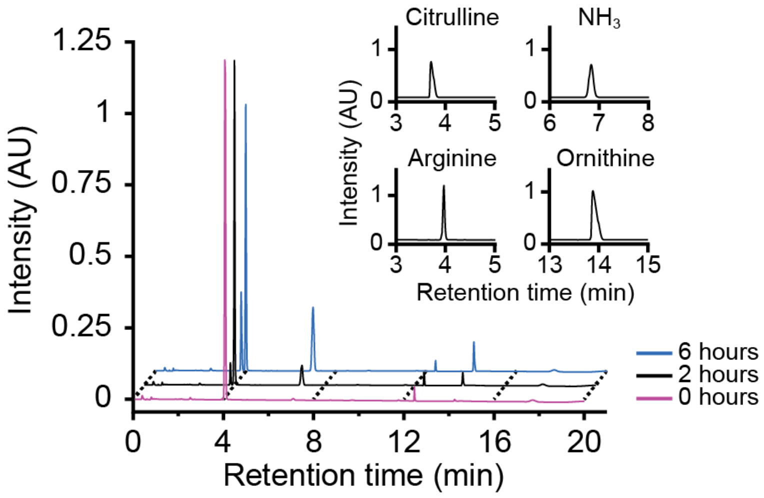

**Figure S7. Raw spectra from HPLC chromatograms**

Representative chromatograms of amino acids and ammonia produced by the vesicles containing the arginine breakdown pathway. Chromatograms were recorded at 269 nm. As expected, at t = 0 only 5 mM arginine is detected (at retention time = 4 min). Citrulline, $NH_3$ and ornithine (at retention time 3.8; 6.8 and 14 min, respectively) appear at the expense of arginine in the chromatograms of samples taken at 2 and 6 hours. Inset: Truncated chromatograms of standard solutions containing 5 mM of a single amino acid or $NH_3$. Such chromatograms were used to determine the retention time and to calibrate the signals.

**Table S5. Enzyme activities for data presented in Fig. 3E**

| Buffer | ArcA | ArcB | ArcC1 |
|---|---|---|---|
| 50 mM KPi | 2.9 ± 0.3 | 231.0 ± 51.5 | 362.5 ± 22.8 |
| 300 mM KPi | 4.8 ± 2.3 | 121.1 ± 49.5 | 103.3 ± 121.3 |

All values are given in µmol * $min^{-1}$ * $mg^{-1}$.
